# Redox stress reshapes carbon fluxes of *Pseudomonas putida* for cytosolic glucose oxidation and NADPH generation

**DOI:** 10.1101/2020.06.13.149542

**Authors:** Pablo I. Nikel, Tobias Fuhrer, Max Chavarría, Alberto Sánchez-Pascuala, Uwe Sauer, Víctor de Lorenzo

## Abstract

The soil bacterium and metabolic engineering platform *Pseudomonas putida* tolerates high levels of endogenous and exogenous oxidative stress, yet the ultimate reason of such property remains unknown. To shed light on this question, NADPH generation routes—the metabolic currency that fuels redox stress responses—were assessed when *P. putida* KT2440 was challenged with H_2_O_2_ as proxy of oxidative conditions. ^13^C-tracer experiments, metabolomics and flux analysis, together with inspection of physiological parameters and measurement of enzymatic activities, revealed a substantial flux reconfiguration under oxidative stress. In particular, periplasmic glucose processing was rerouted to cytoplasmic oxidation, and cyclic operation of the pentose phosphate pathway led to significant NADPH fluxes, exceeding biosynthetic demands by ~50%. This NADPH surplus, in turn, fuelled the glutathione system for H_2_O_2_ reduction. These properties not only contribute to the tolerance of *P. putida* to environmental stresses, but they also highlight the value of this host for harsh biotransformations.

## Introduction

Oxidative stress is an unavoidable consequence of aerobic cell respiration, characterized by the systemic manifestation of reactive oxygen species (ROS) followed by activation of multiple responses deployed to detoxify reactive intermediates^1–4^. Peroxides and free radicals damage virtually all macromolecular components of the cell (e.g. proteins, lipids and DNA^5,6^), and mechanisms to prevent (or repair) the damage exerted by ROS are key components of virtually all biological systems^7^. Long-term transcriptional responses that scavenge ROS, for instance, appear to be conserved across species^8^. Model microorganisms such as *Escherichia coli* and *Saccharomyces cerevisiae* have been shown to activate coordinated transcriptional responses that include up-regulation of ROS-quenching superoxide dismutases, catalases and glutathione/glutaredoxin recycling systems^9–12^. Similarly, mammalian cells display long-term anti-oxidative responses for ROS detoxification and, depending on the severity of oxidative stress, initiate either pro-survival gene expression strategies to support NADPH formation^13^, ROS clearance and (in most cases) DNA repair^14^, or they commit cell death programs^15,16^. However, until transcriptionally-regulated defence mechanisms become fully operational, cell survival depends on the basal activity of the above-mentioned enzymes and availability of antioxidants (e.g. reduced glutathione^17,18^) to scavenge at least part of the ROS generated by respiration or direct oxidative damage. Besides transcriptional responses, metabolic programmes are rapidly deployed upon exposure of bacteria, yeast and mammalian cells to oxidative stress^19–22^, thereby providing reducing power to fuel ROS detoxifying enzymes^23^. However, metabolic mechanisms to counteract oxidative stress in non-model organisms remain poorly studied.

Environmental bacteria are constantly exposed to different types of stress in the niches they colonize. *Pseudomonas putida*, a strictly-aerobic, Gram-negative soil bacterium characterized by a remarkable metabolic versatility^24–27^, is equipped with the enzymatic machinery needed to catabolize both natural and recalcitrant aromatic compounds besides sugars and organic acids^28^. While many of the scenarios where *P. putida* meets such environmental pollutants are oxidative *per se*, metabolic pathways for their biodegradation often involve harsh redox reactions prone to generate ROS endogenously^29^. The adaptability of *P. putida* KT2440 (type strain of this species^30^) to adverse conditions, including exposure to aromatic substrates that cause endogenous oxidative stress, is wired to a unique metabolic architecture. Glucose catabolism in *P. putida* relies on the Entner-Doudoroff (ED) pathway^31^, which starts with 6-phosphogluconate (6PG) as the substrate, formed *via* separate and converging routes for hexose phosphorylation (in the cytoplasm) or oxidation (in the periplasm)^32^. Additionally, the *EDEMP cycle*, a combination of enzymes from the ED, the pentose phosphate (PP) and the (incomplete) Embden-Meyerhof-Parnas (EMP) pathways^33^, processes hexoses phosphate to feed trioses into the lower catabolism, i.e. downwards phosphoenolpyruvate (PEP), pyruvate (Pyr) and the tricarboxylic acid (TCA) cycle. The metabolic network of *P. putida* KT2440 and biochemical reactions considered in this study are summarized in **Supplementary Fig. S1** and **Supplementary Table S1** respectively. In addition to catabolic processing of sugars, the operation of the EDEMP cycle entails redirecting part of the trioses phosphate back to hexoses phosphate *via* the gluconeogenic activity of the EMP branch of the cycle^34^. As such, cyclic sugar catabolism contributes to a slight catabolic overproduction of NADPH in glucose cultures^33^. The EDEMP cycle has been also shown to be important for the hierarchical consumption of sugars and aromatic compounds^35^. While the main sink of NADPH is the anabolic use of reducing power for biomass formation, this redox cofactor is also key to counterfeit oxidative stress since it serves as the reductant for several ROS detoxifying enzymes and mechanisms^36^, e.g. glutathione regeneration. However, the connection between the operation of central carbon metabolism and oxidative stress responses remained elusive thus far.

In this work, the reshaping of the central metabolism in *P. putida* KT2440 in response to sub-lethal doses of oxidative stress elicited by exposure to hydrogen peroxide (H_2_O_2_) has been explored. The experimental strategy encompassed the use ^13^C-tracer experiments, metabolomics and metabolic flux analysis, combined with determination of physiological parameters, *in vitro* measurement of key enzymatic activities and assessment of glutathione-mediated ROS detoxifying mechanisms. The results revealed adaptions in the operation of the EDEMP cycle, concomitant with a decrease in the flux through sugar oxidation pathways for glucose processing, which helps replenishing the NADPH pool upon oxidative challenges. These metabolic adjustments are discussed in the light of the lifestyle of *P. putida* (and related environmental bacteria) and the value of this species as robust platform for hosting redox-demanding biotransformations.

## Results

### Quantitative physiology analysis of *P. putida* exposed to sub-lethal doses of oxidative stress

In order to capture the metabolic effects of oxidative stress in *P. putida* KT2440 without compromising cell viability, the experimental design (summarized in **Fig. 1a**) entailed exposure of the cultures, grown in M9 minimal medium with glucose until mid-exponential phase [optical density at 600 nm (OD_600_) of ca. 0.5], to H_2_O_2_ at 1.5 mM. Cultures with different ^13^C-tracers ([1-^13^C_1_]-, [6-^13^C_1_]- and [U-^13^C_6_]-glucose, all of them added at 20 mM) were grown in parallel to resolve the relative contribution of cyclic EDEMP, PP and ED pathway to sugar processing. At 1.5 h after the oxidative stress treatment, samples were collected to assess quantitative physiology parameters, flux ratio analysis and to perform *in vitro* biochemical assays. This experimental setup resulted in similar growth patterns in the absence and presence of the oxidative stress agent (**Fig. 1b**), and neither the specific growth rate (μ) nor the final cell density (plateauing at ca. 10 h of cultivation) were significantly affected by the addition of H_2_O_2_ to the cultures. Quantitative analysis of physiology parameters confirmed that this was the case (**Fig. 1c**), with μ, the specific rate of glucose uptake (*q*_S_) and the yield of biomass on substrate (*Y*_X/S_) being essentially the same in control and H_2_O_2_–treated experiments. The formation and secretion of organic acids to the extracellular milieu, a typical feature of glucose-grown *P. putida*, decreased significantly in the presence of the stressor. *P. putida* typically oxidizes glucose in the periplasm into gluconate and 2-ketogluconate^31^, which are then taken up by the cell^37^. The about halved concentrations of the two products of sugar oxidation in H_2_O_2_–treated cultures indicates a decrease in periplasmic oxidation upon exposure to stressful conditions. The concentration of both acids peaked during exponential growth, and the corresponding molar yields of gluconate and 2-ketogluconate on glucose in control experiments were *y*_G/S_ = 0.31 ± 0.09 and *y*_K/S_ = 0.12 ± 0.02 C-mol C-mol^−1^, respectively. In the presence of oxidative stress, *y*_G/S_ and *y*_K/S_ were 0.19 ± 0.08 and 0.05 ± 0.01 C-mol C-mol^−1^, suggestive, again, of a redistribution of metabolic fluxes between oxidative (i.e. glucose → gluconate → 2-ketogluconate) or phosphorylative [i.e. glucose → glucose-6-phosphate (G6P)] processing routes of the carbon source (**Supplementary Fig. S1**). Since a shift from periplasmic to cytoplasmic glucose oxidation would entail higher NADPH formation, we next looked for direct evidence of increased intracellular fluxes through these pathways.

**Figure 1.**
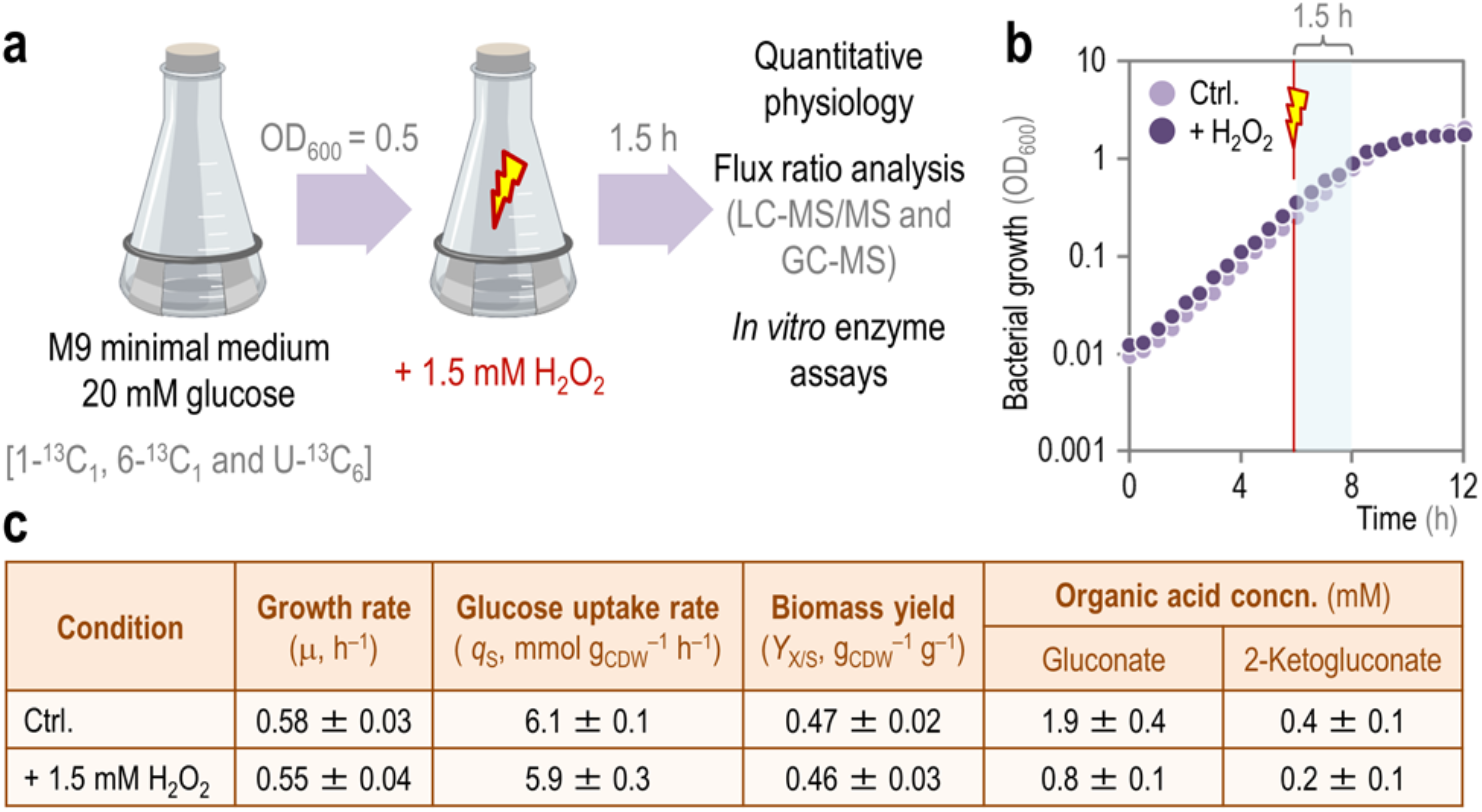
Experimental overview and quantitative physiology parameters for *P. putida* KT2440 cultures. **a.** Overview of the experimental design. All experiments were conducted at least in biological triplicates. OD_600_, optical density measured at 600 nm. **b.** Typical growth curves for control (Ctrl.) experiments and H_2_O_2_-stressed cultures. Note that samples were taken 1.5 h after addition of H_2_O_2_. **c.** Physiological parameters of *P. putida* KT2440 batch cultures. All values represent the mean at least three biological replicates. The specific growth rate (μ) and the specific rate of carbon uptake (*q*_S_) were determined during exponential growth by linear regression of log-transformed OD data. The yield of biomass on substrate (*Y*_X/S_) was calculated at 24 h, after glucose was completely exhausted. The extracellular concentration (concn.) of organic acids reported is the maximal reached during the whole culture period. CDW, cell dry weight.

### PP pathway metabolite pools increase upon exposure of *P. putida* to oxidative stress

To assess the adaptions within the metabolic network of *P. putida* upon exposure to oxidative stress, the relative abundance of key metabolites was quantified by LC-MS/MS. Out of the metabolites measured, the species that showed the most significant changes in terms of abundance were intermediates of the PP pathway, the EMP route and the TCA cycle—albeit in opposite directions (**Fig. 2a**). In particular, the relative abundance of 6PG (key metabolic node where oxidative and phosphorylative pathways for glucose processing converge; **Supplementary Fig. S1**), ribose-5-phosphate (R5P), ribulose-5-phosphate (Ru5P), xylulose-5-phosphate (Xu5P) and sedoheptulose-7-phosphate (S7P) augmented significantly in H_2_O_2_–stressed cells, ranging from a 2- (Xu5P) to 7.6-fold (Ru5P) increase. At the same time, the relative abundance of glycolytic intermediates of the EMP route decreased roughly by half (e.g. fructose-6-phosphate, F6P) and up to 70% (e.g. Pyr) when cells were exposed to oxidative stress. Intermediates of the TCA cycle shared the same fate, and their relative abundance was reduced at least by 60% (e.g. isocitrate, ICT) and up to ca. 90% (e.g. 2-ketoglutarate, KG; succinate, SUC; and malate, MAL) in stressed cells. The changes in the relative abundance of other metabolites in the biochemical network of *P. putida* KT2440 upon exposure to the oxidative stress agent are listed in **Supplementary Data 1**. Taken together, these results indicate increased fluxes through the PP pathway coupled to a decrease in the catabolic activities of the EMP route and the TCA cycle during oxidative stress. However suggestive, these metabolomic data cannot be used to infer relative or net fluxes, and flux ratio analysis was employed to piece together the contribution of fluxes within the upper metabolism of *P. putida* to the pool of glycolytic intermediates.

**Figure 2.**
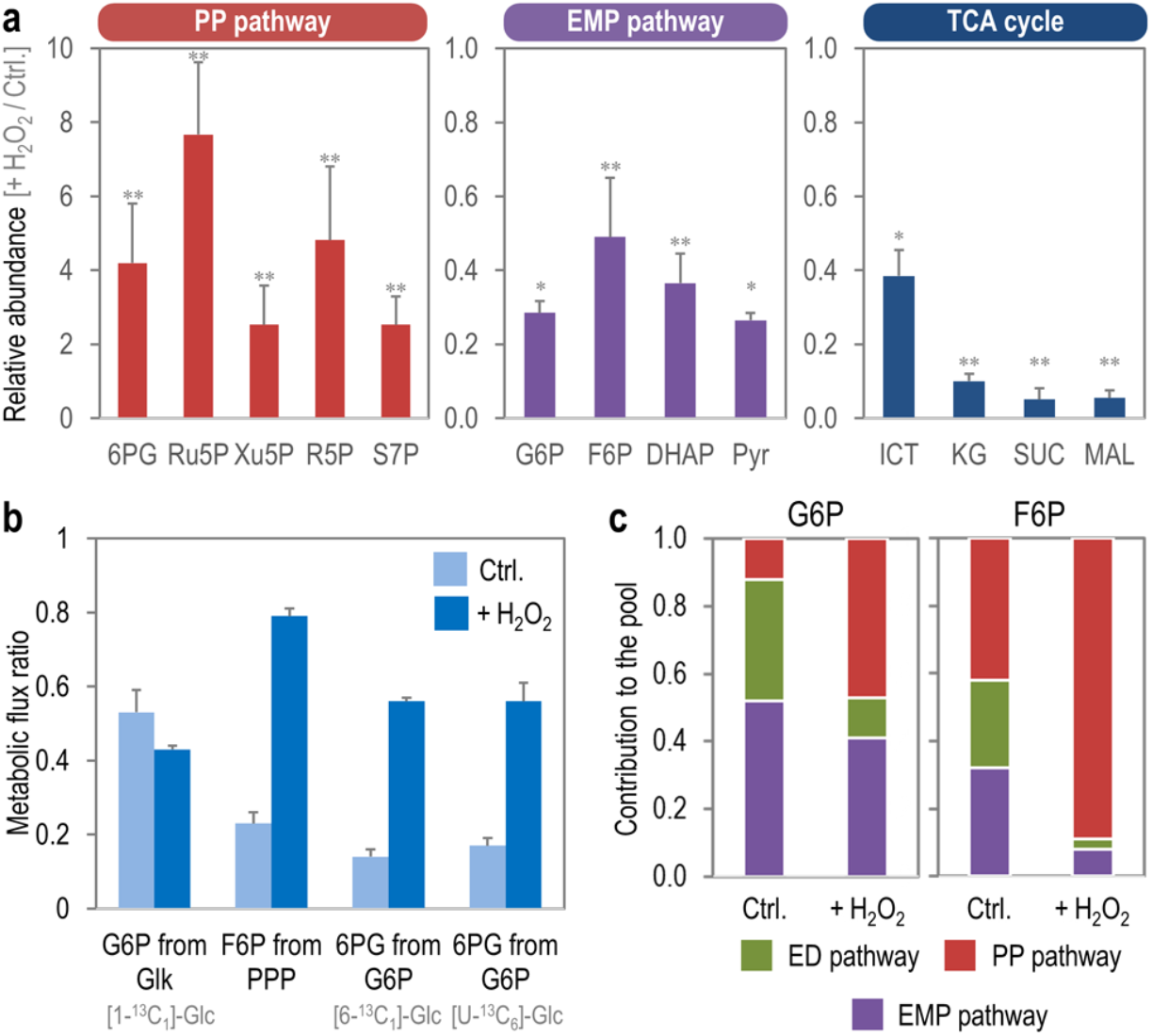
Metabolite levels in *P. putida* KT2440 under oxidative stress. **a.** Relative abundance of selected metabolites, grouped according to the biochemical block they belong to (i.e. PP pathway, EMP pathway and TCA cycle). Relative metabolite abundance is expressed as the ratio between H_2_O_2_–induced oxidative stress and control (Ctrl.) conditions, derived from summed ion abundance of all isotopes (counts). Data from experiments in the presence of [1-^13^C_1_]- and [6-^13^C_1_]-glucose (Glc.) are averaged. Bars represent the mean value of the abundance ± standard deviations of triplicate measurements from at least two independent experiments. Single (*) and double asterisks (**) identify significant differences at *P* < 0.05 and *P* < 0.01, respectively, as assessed by the Student’s *t* test. **b.** Changes in selected metabolic flux ratios in upper metabolism upon exposure of the cells to H_2_O_2_. The ^13^C-labeled substrate used in each experiment is indicated. Bars represent averages from three independent experiments, and standard deviations were calculated using the covariance matrices of the respective mass distribution vectors by applying the Gaussian law of error propagation. **c.** Relative pathway contribution to the G6P and F6P pools. The contribution of each of the metabolic pathways to the sugar phosphate pool under oxidative stress conditions is indicated with different colours. All abbreviations in this figure are as indicated in the legend to **Fig. S1** in the Supplementary Information. CDW, cell dry weight.

### Increased cyclic operation of PP but not ED pathway upon exposure to oxidative stress

The relative contributions of reverse (i.e. gluconeogenic) flux from triose phosphates or through the PP pathway to the hexose phosphate pool was elucidated by using relative flux ratios derived from [1-^13^C_1_]- and [6-^13^C_1_]-glucose–labeling experiments. Calculation of relative ratios around the hexose node was possible given that the reaction catalyzed by 6-phosphofructo-1-kinase is absent in *P. putida*^38,39^, and by considering the reactions mediated by GntZ (6PG dehydrogenase) and Edd (6PG dehydratase) to be irreversible. In particular, using [6-^13^C_1_]-glucose as a tracer enables the resolution of fluxes from the cyclic PP and ED pathways back to hexose phosphates because the C6 position is maintained in both routes and can lead to double-labeled hexose phosphate molecules. Flux ratio analysis supported the notion that fluxes through the PP pathway significantly responded to H_2_O_2_ addition (**Fig. 2b**), in particular in the oxidative branch of this metabolic pathway.

Experiments conducted in the presence of [1-^13^C_1_]-glucose suggested that the contribution of glucose kinase (Glk, *v*_3_ in **Supplementary Fig. S1**) to the G6P pool was not significantly affected by oxidative stress, whereas the fraction of 6PG from G6P *via* G6P dehydrogenase (Zwf, *v*_7_ in **Fig. S1**) increased by at least 3.5-fold (as deduced from experiments conducted with either [6-^13^C_1_]- or [U-^13^C_6_]-glucose) in stressed *P. putida*. The increase was even more pronounced for the flux ratio reflecting the fraction of F6P from the PP pathway (4.5-fold higher in cultures treated with H_2_O_2_ than in control experiments). The full list of flux ratios, including those for reactions in lower catabolism, is presented in **Supplementary Table S2**, and the GC-MS data are given in **Supplementary Data 2**. These results provided further evidence for a flux rerouting from periplasmic glucose oxidation to the NADPH-generating PP pathway as a direct metabolic response to oxidative stress. We next quantified net fluxes as the basis for accurate estimation of NADPH production in response to oxidative stress.

### Redistribution of net fluxes in *P. putida* as a response to sub-lethal oxidative stress

Combining the quantitative physiology data (**Fig. 1c**) with the flux ratios as additional constraints (**Fig. 2b** and **Supplementary Table S2**), we estimated net fluxes through the network (**Fig. 3**, and **Supplementary Table S3**). Since the relative contribution of the direct phosphorylation of gluconate to 6PG and that of the indirect (NADPH-dependent) route though 2-ketogluconate could not be resolved from ^13^C-labeling data, we determined the *in vitro* enzyme activities of gluconate kinase (GnuK, i.e. direct phosphorylation of the sugar) and 2-ketogluconate-6-phosphate reductase (KguD, i.e. 6PG from 2-ketogluconate) as a proxy. GnuK and KguD activities in cell-free extracts of *P. putida* KT2440 were similar under control and stress conditions, and we thus used a GnuK/KguD activity ratio of 0.87/0.13 to constrain the flux split at the gluconate branch point.

**Figure 3.**
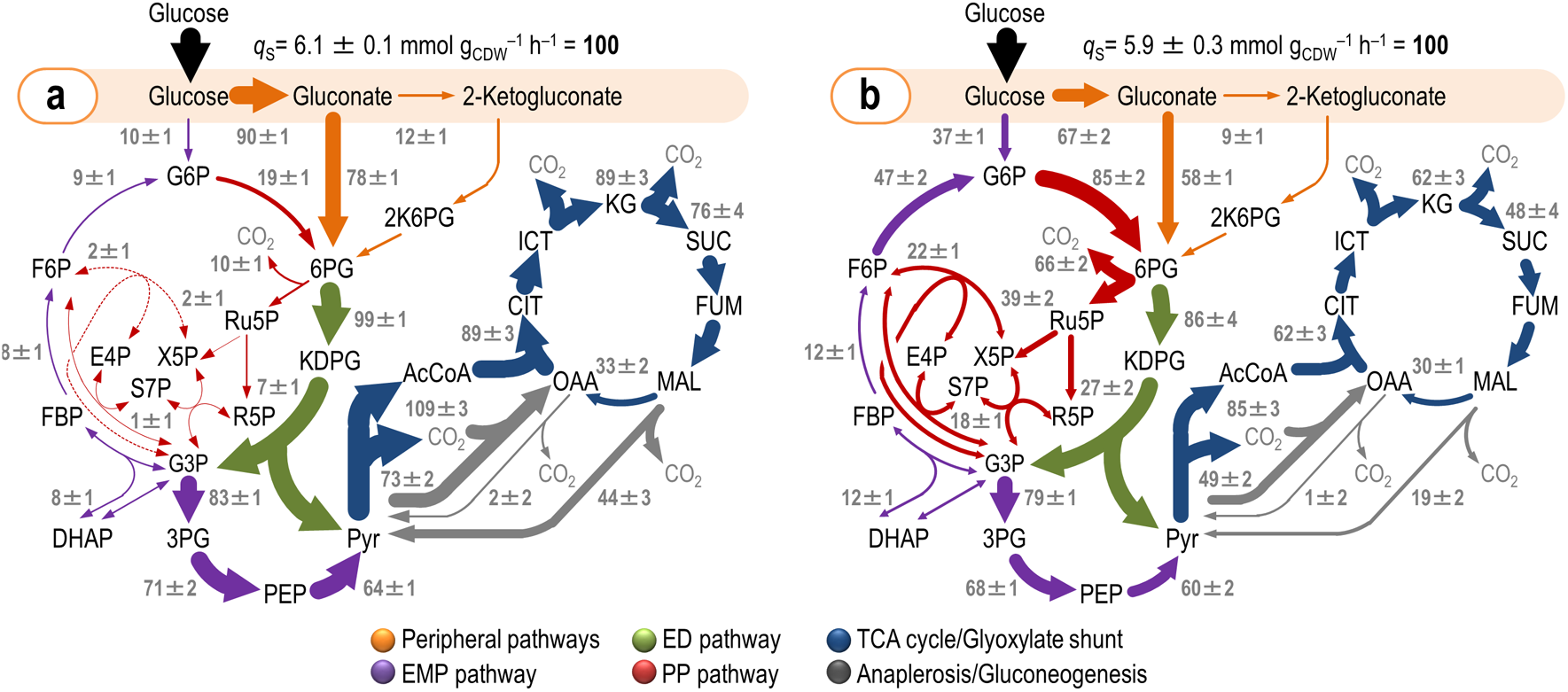
*In vivo* carbon flux distribution in glucose-grown *P. putida* KT2440 obtained from ratio-constrained flux balance analysis. All fluxes, calculated under control conditions (**a**) and in the presence of H_2_O_2_–induced oxidative stress (**b**), were normalized to the specific glucose uptake rate (*q*_S_), and the width of each arrow is scaled to the relative flux. Dashed lines indicate that no significant flux through the corresponding biochemical step was detected under the conditions tested. Abbreviations used for the metabolic intermediates and the main metabolic blocks within the biochemical network are given in the legend to **Fig. S1** in the Supplementary Information. CDW, cell dry weight.

Under control conditions, glucose was processed mostly through the periplasmic pathway with about 90% of the sugar uptake rate channeled *via* Gcd (**Fig. 3a**). Most of the gluconate fed the 6PG node, with a very high flux through the ED route and a ca. 10% recycling of trioses phosphate through the EDEMP cycle. Upon exposure to H_2_O_2_, *P. putida* substantially re-routed its carbon fluxes (**Fig. 3b**). Periplasmic glucose oxidation decreased, and instead much more glucose was imported into the cytoplasm and converted to 6PG through the NADP^+^-dependent Zwf. Since the glucose uptake rate was essentially identical in the two experimental conditions, this large relative flux change reflects a likewise large change in absolute fluxes. In particular, a major change in flux distribution was detected in the non-oxidative branch of the PP pathway. Exposure of the cells to H_2_O_2_ increased the GntZ flux by more than 6-fold, which fed the pools of other pentoses phosphate (as suggested by the measurement of their relative abundance increase, **Fig. 2**). The backward flux in this route fed the F6P and T3P node, resulting in carbon recycling *via* glucose-6-phosphate isomerase (Pgi, in the gluconeogenic F6P → G6P direction) and Zwf (G6P → 6PG). Accordingly, fluxes through Pgi and Zwf in H_2_O_2_–treated cells increased by 5.2- and 4.5-fold, respectively. The ED route was still the predominant catabolic pathway for 6PG under oxidative stress, albeit with a slight decrease in the flux values as compared to control conditions. Gluconeogenic fluxes *via* fructose-1,6-bisphosphate aldolase (Fda) and fructose-1,6-bisphosphatase (Fbp) had a small increase (1.5-fold) in cultures exposed to oxidative stress, and flux through the lower, catabolic branch of the EMP pathway (from glyceraldehyde-3-phosphate to PEP and Pyr) was reduced by ca. 10% under oxidative stress. The diminished input of the ED and the EMP catabolism under oxidative stress propagated into a relatively low flux towards acetyl-CoA (i.e. Pyr dehydrogenase) and fluxes within the TCA cycle. No significant activity of the glyoxylate shunt could be detected under any of the experimental conditions tested. Anaplerotic routes within the Pyr shunt, typical of glucose-grown *P. putida*, had a likewise lower activity in H_2_O_2_-stressed cells. The impact of oxidative stress-mediated re-arrangement of carbon fluxes on redox metabolism was investigated next.

### Re-arrangement of net fluxes in *P. putida* under oxidative stress increases NADPH supply

Considering that the flux through Zwf and GntZ, two of the major NADP^+^-dependent dehydrogenases in the biochemical network, showed a significant increase under oxidative stress, the overall net NADPH production increased substantially. This reconfiguration presumably matches the increased NADPH demand to regenerate glutathione (and other anti-oxidants) for H_2_O_2_ reduction. To assess this impact quantitatively, the net rate of NADPH production (*R*_NADPH_) was calculated according to ∑_i_ (*r*_i_^F^ – *r*_i_^C^), where *r*_i_^F^ and *r*_i_^C^ represent the rates of NADPH formation and consumption, respectively, of all major dehydrogenases potentially carrying significant flux in the biochemical network (see **Supplementary Fig. S1**). Zwf, GntZ, Gap, Icd, Mdh and MaeB were considered as potential inputs to NADPH formation for this analysis, whereas biomass formation and the KguD activity were included as NADPH sinks. The values of *r*_i_ were derived from the actual previously calculated net fluxes, and the cofactor specificity of all dehydrogenases under both saturating, *in vitro* conditions and non-saturating, *quasi in vivo* conditions (**Supplementary Table S4**) was used to derive the *r*_i_ values as previously indicated. By combining these parameters, *R*_NADPH_ was estimated for *P. putida* in both control and H_2_O_2_-treated experiments (**Fig. 4**). When using cofactor specificities determined under saturating, *in vitro* conditions, NADPH formation and consumption in control experiments added up (i.e. *R*_NADPH_ = 0, **Fig. 4a**), whereas adjusting these specificities according to non-saturating, *quasi in vivo* determinations revealed a slight catabolic overproduction of reducing power characteristic of strain KT2440 (*R*_NADPH_ = 1.3 mmol g_CDW_^−1^ h^−1^, **Fig. 4b**). In H_2_O_2_-stressed cells, *R*_NADPH_ was > 0 irrespective of the cofactor specificity used to calculate the individual *r*_i_ values (**Figs. 4c** and **4d**). In particular, when applying the specificity coefficients derived from non-saturating, more realistic *quasi in vivo* determinations, *R*_NADPH_ increased by 3.6-fold (**Fig. 4d**) as compared to the control condition (**Fig. 4b**). Taken together, these results expose a ca. 50% surplus in NADPH formation under oxidative stress conditions that becomes available to counteract ROS accumulation.

**Figure 4.**
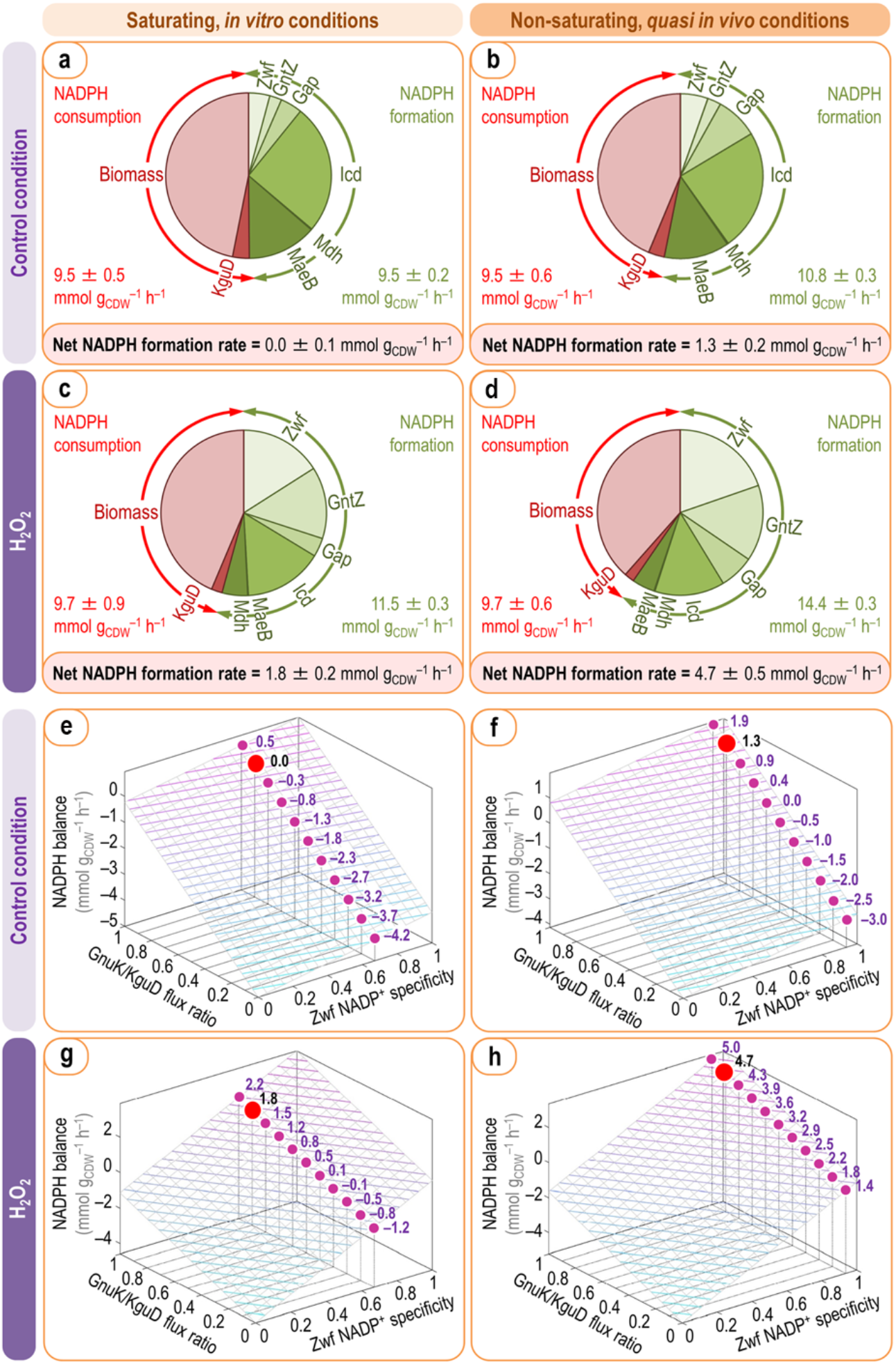
Dynamic NADPH balance in *P. putida* KT2440 upon oxidative stress. Overall NADPH balances and sensitivity analysis for the wild-type strain KT2440 under control (**a**, **b**, **e** and **f**) or H_2_O_2_–induced oxidative stress conditions (**c**, **d**, **g** and **h**) using cofactor specificities of major dehydrogenases under saturating conditions (left column) or *quasi in vivo* conditions (right column). NADPH formation was determined from carbon fluxes through redox cofactor-dependent reactions (**Fig. 3** and **Table S1** in the Supplementary Information) multiplied by experimentally-determined relative cofactor specificities. NADPH consumption was calculated from requirements for biomass production and KguD activity. NADPH formation (green) and consumption (red) rates are individually indicated (**a**–**d**). Dependence of the overall rates of NADPH turnover on the relative GnuK/KguD flux ratio and the cofactor specificity of Zwf derived from saturating conditions or *quasi in vivo* conditions (**e**–**h**). Selected values for the net NADPH balance are given for the experimentally-determined NADP^+^ specificity of Zwf (0.937), and the calculated net rate for each conditions is indicated with a red dot. CDW, cell dry weight.

To ensure that our NADPH balance estimates are not biased by key assumptions, we assessed the sensitivity of net NADPH formation as a function of the GnuK/KguD activity ratio at the gluconate node and the NADP^+^-specificity of the dehydrogenase reaction catalyzed by Zwf. The Zwf activity of *P. putida* KT2440 is represented by three isozymes (i.e. Zwf, ZwfA and ZwfB; **Supplementary Table S1**). Even when we have determined NADP^+^ and NAD^+^ specificities for the overall Zwf reaction with in vitro assays, it is likely that the individual isozymes may have different cofactor preferences, which would have a direct effect on the overall redox balance if expressed differently. Irrespective of the cofactor preferences (0.672 and 0.937 either under saturating conditions or *quasi in vivo* conditions, respectively; see **Supplementary Table S4**), *R*_NADPH_ was relatively more responsive to changes in the GnuK/KguD flux ratio in control, non-stressed experiments (**Figs. 4e** and **4f**). The opposite was true for H_2_O_2_–treated cells, with a strong dependence of *R*_NADPH_ on the output of the Zwf flux (**Figs. 4g** and **4h**) mainly because the relative flux through Zwf increased in comparison to the control experiment. These calculations demonstrate that the NADPH surplus observed in stressed cells largely stems from the NADP+-specificity of Zwf, with the GnuK/KguD flux ratio playing a minor role on *R*_NADPH_ values.

### Altered expression of enzymes in glucose catabolism during oxidative stress

In agreement with the relative flux distribution (**Fig. 3**), there was a significant decrease in the activities of enzymes within the peripheral reactions for glucose oxidation under oxidative stress conditions (**Fig. 5a**). The observed decrease was particularly significant in the case of the specific Gad activity (gluconate → 2-ketogluconate), which was roughly halved in cells exposed to H_2_O_2_. Hexose kinase activity, accordingly, increased by 2-fold. The activity of the two NADPH producing dehydrogenases of the PP pathway was more significant, with the specific Zwf and GntZ activities increasing by 4.7- and 9.2-fold, respectively. The *in vitro* activities of two enzymes of the ED pathway, in contrast, remained unchanged within the experimental error in control and H_2_O_2_-stressed cells. All these results are consistent with the previous observations underline the importance of PP pathway for NADPH production when cells are challenged with an oxidative stress agent. We next explored the connection between catabolic NADPH overproduction and mechanisms that counteract oxidative stress such as the glutathione-dependent redox system.

**Figure 5.**
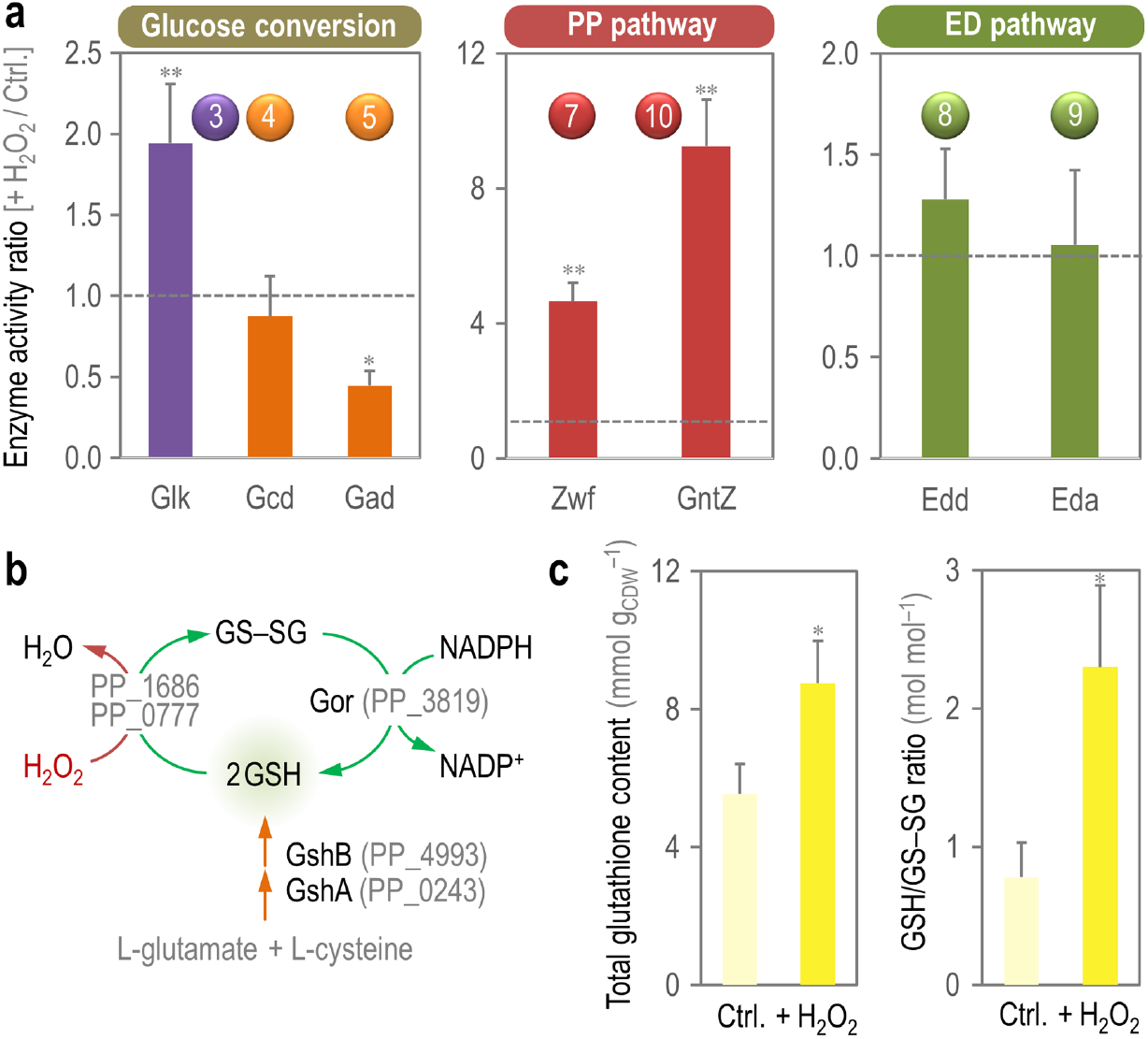
*In vitro* analysis of key enzymatic activities and glutathione metabolism. **a.** Enzyme activity ratios were calculated from the specific activity for each of the indicated reactions assessed under H_2_O_2_–induced oxidative stress and control (Ctrl.) conditions. Each bar represents the mean value of the corresponding ratio ± standard deviations of triplicate measurements from at least two independent experiments, and the horizontal dashed line indicates a ratio = 1 (i.e. no changes in enzymatic activities across conditions). Single (*) and double asterisks (**) identify significant differences at *P* < 0.05 and *P* < 0.01, respectively, as assessed by the Student’s *t* test. Circled numbers identify the enzymes in the biochemical network of **Fig. S1** in the Supplementary Information. **b.** Glutathione metabolism in *P. putida* KT2440. The key activities involved in biosynthesis and recycling of the reduced (GSH) and oxidized (GS–SG) forms of glutathione are indicated along with the corresponding PP identifiers. **c.** Enzymatic determination of total glutathione and the fraction of the oxidized and reduced form. Bars represent the mean value of the corresponding parameter ± standard deviations of duplicate measurements from at least three independent experiments, and the asterisk (*) identifies significant differences between stressed cells and control conditions at *P* < 0.05 as assessed by the Student’s *t* test. CDW, cell dry weight.

### The glutathione system of *P. putida* links H_2_O_2_-induced overproduction of NADPH to ROS quenching

Thiols play several roles in bacteria, including maintenance of the redox balance, fighting ROS and nitrogen reactive species, and detoxification of other toxins and stress-inducing factors. In most organisms, the major thiol is the tripeptide glutathione (GSH, γ–L-glutamyl-L-cysteinylglycine). The formation of GSH from L-glutamate and L-cysteine in strain KT2440 is catalyzed by GshAB (glutamate-cysteine ligase and glutathione synthase, respectively; **Fig. 5b**). The glutathione cycle, connecting the reduced (GSH) and the oxidized (GS–SG) forms of the thiol, acts as a first-line reductant of ROS (**Fig. 5b**) *via* glutathione peroxidases (e.g. PP_0777 and PP_1686) and glutaredoxins. Glutathione reductase (Gor, PP_3819) then replenishes the GSH pool by using NADPH as the reducing currency. Since the glutathione system of *P. putida* has not been explored thus far from a biochemical point of view, the cellular content of both GSH and GS–SG was assessed by means of a biochemical assay in both control and H_2_O_2_-stressed cultures (**Fig. 5c**). Cells exposed to H_2_O_2_ had a 1.6-fold higher total content of glutathione (i.e. GSH and GS–SG) than control experiments, and the composition of the total pool changed significantly upon exposure to oxidative stress. The GSH/GS–SG molar ratio, which reflects the fraction of reduced thiol in the cells, increased by 3-fold in H_2_O_2_–stressed *P. putida*. This observation suggests that the reduced form of glutathione can be sufficiently replenished by the cellular NADPH sources upon an oxidative stress—directly linking the increased NADPH turnover with the pool of antioxidants as known from well-studied species.

## Discussion

Environmental microorganisms are constantly exposed to a variety of physicochemical perturbations that require dynamic and concerted responses to ensure cell survival^40^. In this context, *P. putida* KT2440 provides a unique experimental system to explore the interplay between stress and metabolic adaption. This understanding has not only a fundamental biological interest, but also a practical one in view of the growing demand of robust *chassis* for synthetic biology and metabolic engineering^41^. In this sense, the results of this study strengthen the notion that network-wide adaption of metabolic fluxes^42,43^ accompanies (and plausibly, enables) what has been traditionally described—mostly at a genetic and regulatory level—as *general stress response*^44–47^. Metabolic adaption to oxidative stress has been recently revisited by Christodoulou *et al.*^23^, demonstrating how reserve flux capacity in the PP pathway empowers *E. coli* to rapidly respond to H_2_O_2_ by increased flux through Zwf and Gnd. It was postulated that inhibition of Zwf caused by NADPH (either by a competitive, allosteric or combined mechanism) is lifted under these conditions—releasing ‘reserve’ flux through the oxidative PP pathway. Moreover, the activity of glyceraldehyde-3-phosphate is lowered due to ROS-mediated damage, which causes a metabolic *jam* in the metabolite pools of the PP route as previously shown in yeast^48^. Product inhibition of Zwf is lowered due to NADPH drainage to counteract oxidative stress^49^, and the sudden enlargement of the 6PG pool is channelled *via* Gnd. Recently, such an NADPH replenishing mechanism catalysed by Zwf and Gnd has been recognized as a general strategy in enteric bacteria, yeast and mammalian cells^50^. Here, we asked how *P. putida*, which does not have a linear, EMP-based glycolysis and processes glucose mostly through the conversion of the sugar into gluconate, reacts to oxidative stress. Since glycolysis is not an alternative glucose degradation route and therefore *P. putida* cannot alter the split ratio between the EMP and PP pathways, we hypothesized that the EDEMP cycle could contribute to increased NADPH production under oxidative stress *via* the PP pathway branch.

Metabolite changes and flux analysis revealed that the lower (EMP) glycolysis and the PP pathway were affected by H_2_O_2_ as compared with untreated controls. In a simplified model of the upper pathways for sugar processing (**Fig. 6a**), the output of NADPH is largely governed by the fluxes through Zwf and GntZ—that, in turn, respond to the split between oxidative conversion of glucose into gluconate or phosphorylation into G6P, and the recycling activity of the EDEMP cycle. As such, the overall NADPH turnover is dependent on *r_s_* (**Fig. 6b**), and is further modulated by the fraction of glucose oxidized in the bacterial periplasm. Consequently, a low activity of the EDEMP cycle contributes to NADPH formation when cells grow in the absence of any oxidative stress. Upon an oxidative challenge, the backward flux from the PP and ED routes to F6P increases >3 fold, and the ED flux through lower glycolysis decreases due to re-arrangement. The jump in the activities through the cyclic PP pathway is most remarkable, as *P. putida* maintains very low fluxes within this metabolic block under normal growth conditions^51^. Such adaption mechanism entails a significant increase in the activities of enzymes needed for a cyclic operation that burn carbon into CO_2_ to generate NADPH. The overall biomass yield is nevertheless similar to that under control conditions because CO_2_ production by cellular respiration (i.e. pyruvate dehydrogenase and TCA cycle) is reduced. Remarkably, *P. putida* deploys a survival strategy that differs from that observed in the close relative species *P. fluorescens*, where production of α-ketoacids and ATP formation prevails under oxidative stress *via* several converging metabolic mechanisms^46^.

**Figure 6.**
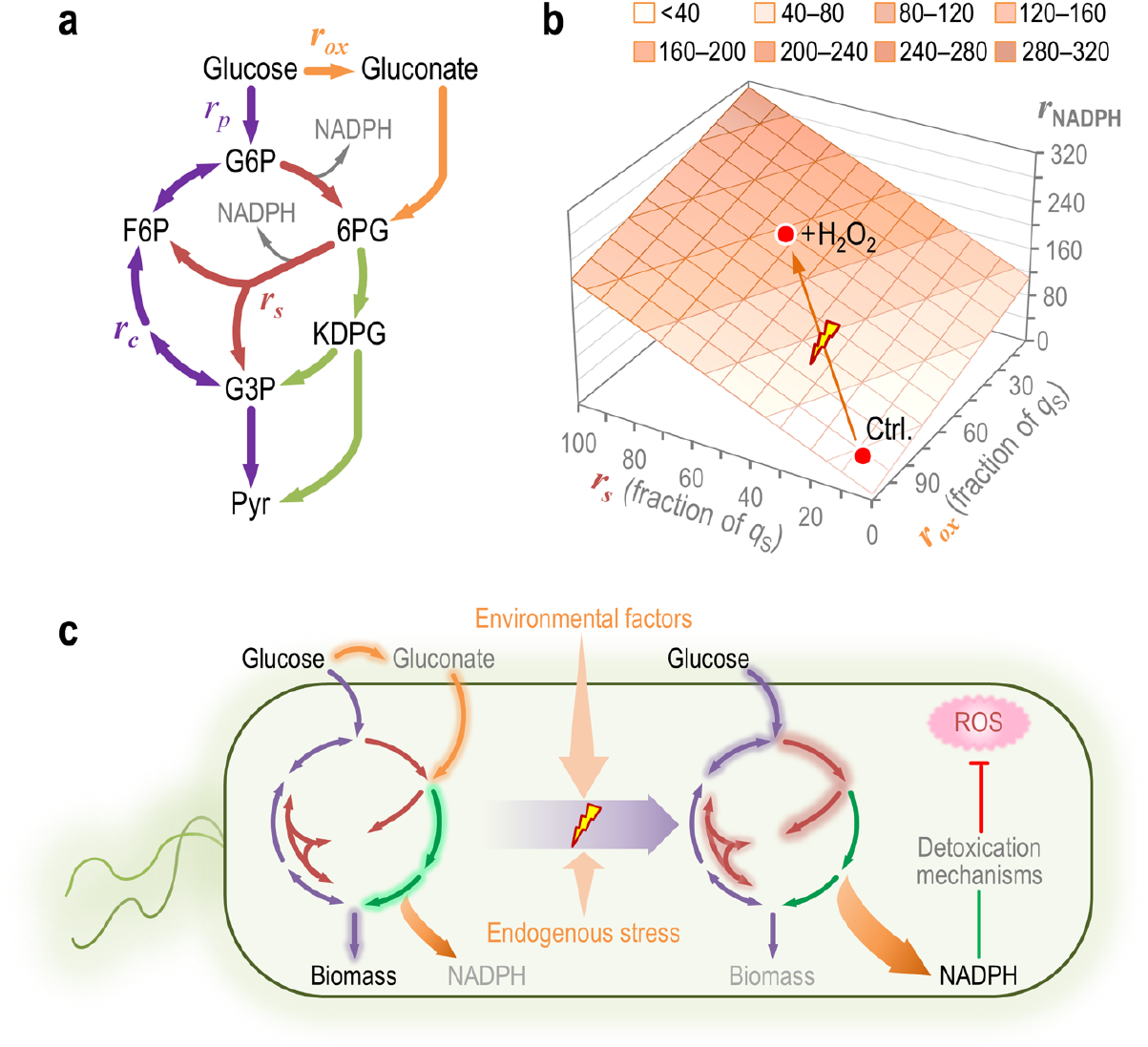
The architecture of central carbon metabolism in *P. putida* enables rapid supply of NADPH upon oxidative stress. **a.** Schematic representation of the upper metabolism of *P. putida* KT2440. Several biochemical reaction have been lumped to illustrate the main routes for carbon circulation (see **Fig. S1** in the Supplementary Information for details and abbreviations). Note that the total rate of carbon uptake (*q*_S_) is split between glucose phosphorylation and oxidation to gluconate, such that *q*_S_ = *r_p_* + *r_ox_*. The overall cycling flux of trioses phosphate towards hexoses phosphate is indicated as *r_c_* and the flux through the PP pathway shunt is termed *r_s_*. Cofactors other than NADPH have been omitted in the drawing for the sake of clarity. **b.** Functional relationship between *R*_NADPH_, the rate of NADPH formation within the simplified metabolic network of panel **a**, and the fluxes through the oxidative loop for glucose processing and the PP pathway shunt. All (arbitrary) values are given as a fraction of *q*_S_, and the experimental conditions tested in this work are indicated with red dots (Ctrl., control conditions). **c.** General model for flux distribution in the upper metabolic domain of *P. putida*. Under normal growth conditions, glucose is processed mostly through its oxidative conversion to gluconate, and the EDEMP cycle provides intermediates for biomass, with a very low flux through the PP pathway. Upon oxidative stress conditions (exerted either by endogenous or external perturbations), a rapid increase of fluxes *via* the PP pathway shunt replenishes the intermediates within upper metabolism and provides a direct source of NADPH that can be coupled to anti-oxidant defence mechanisms against reactive oxygen species (ROS). Note that fluxes predominant under each condition are highlighted.

Increasing evidence points to glutathione peroxidase as a key survival mechanism in stressed bacteria^52–54^—and *P. putida* seems to be no exception. Glutathione peroxidase-dependent reduction of ROS requires a continuous supply of NADPH for regeneration^55^. Upon oxidative stress, the glutathione-based detoxification of ROS drains a lot of NADPH that must be replenished *via* the mechanisms explained above. Importantly, the results of this study reflect a situation where the immediate, early metabolic response (which in *E. coli* takes place within the first 10 s after exposure to the stress agent) is blended with early transcriptional responses (e.g. activation of inducible defence mechanisms, which take several minutes to occur). Experimental evidence indicates that there is a multi-layered array of defence strategies against oxidative stress in *P. putida*, including (i) several catalases and superoxide dismutases^56,57^, (ii) ferredoxin-NADP^+^ oxido-reductases^58^, (iii) polyphosphate-dependent NAD^+^ and NADH kinases^59^ and (iv) stress-induced transhydrogenation activities^60^. However important, these mechanisms can be classified as (relatively) late responses to oxidative insults as they mostly depend on transcriptional regulation, and the re-arrangement of metabolic fluxes to ensure NADPH supply appears to represent a first line of defence as indicated in other organisms^61^.

Finally, the data presented in this work highlights the value of *P. putida* as a *chassis* for synthetic biology-inspired metabolic engineering. In one hand, the flexibility to reconfigure fluxes that supply NADPH helps counteracting ROS resulting from harsh redox reactions to become excessively mutagenic, increasing genetic and genomic stability of engineered catabolic modules^62^. On the other hand, stress-quenching mechanisms enable *P. putida* to sustain diversification of catabolic pathways (e.g. for novel aromatic substrates) involving interim, faulty reactions that create ROS due to substrate misrecognition^63^. In either circumstance, an abundant, conditional NADPH supply of is a key feature to ensure cell survival and durability in the scenarios indicated herein. Together with the capability of *P. putida* KT2440 to adapt to stressful, industrial-scale conditions^64^, these properties are being exploited to enlarge the panoply of microbial platforms for industrial and environmental whole-cell biocatalysis.

## Methods

### Bacterial strain and culture conditions

Wild-type *P. putida* strain KT2440 was used throughout this study^65^, and it was routinely grown in LB medium^66^. Quantitative physiology experiments were carried out in M9 minimal medium containing 6 g L^−1^ Na_2_HPO_4_, 3 g L^−1^ KH_2_PO_4_, 1.4 g L^−1^ (NH_4_)_2_SO_4_, 0.5 g L^−1^ NaCl, 0.2 g L^−1^ MgSO_4_·7H_2_O and 2.5 mL L^−1^ of a trace elements solution^67^. Either natural or ^13^C labelled glucose at 20 mM was used as the sole carbon source. Growth was estimated by measuring the OD_600_ after diluting the culture with 9 g L^−1^ NaCl. Correlation factors between cell dry weight (CDW) and OD_600_ were determined in batch cultures^68^. All cultures were started with an isolated colony from an LB medium plate, suspended in 5 mL of M9 minimal medium in a test tube. After an 18-h incubation, this culture was used to inoculate fresh medium at an OD_600_ of 0.01. Working cultures were cultivated in 250-mL Erlenmeyer flasks containing medium up to one-fifth of their nominal volume. H_2_O_2_ was used as an oxidative stress agent at 1.5 mM, added at OD_600_ = 0.5 (mid-exponential phase). Cultures were incubated for an additional 1.5 h, and samples were taken for analyses as explained below.

### Determination of physiological parameters

Regression analysis was applied during exponential growth to calculate [i] maximum specific growth rate (μ), [ii] biomass yield on substrate (*Y*_X/S_), [iii] specific rate of glucose consumption (*q*_S_), and [iv] molar yield of organic acids on glucose (*y*_P/S_). CDW was quantified by harvesting cells by fast filtration in pre-weighed nitrocellulose filters (0.45 μm), washed twice with 9 g L^−1^ NaCl, and dried at 105°C to a constant weight. Glucose was assayed using a commercial kit from R-Biopharm AG (Darmstadt, Germany). Gluconate and 2-ketogluconate were measured in culture supernatants as described previously^60^.

### Determination of relative metabolite concentrations and ^13^C-labeling patterns by LC-MS/MS

Cultures were grown on either 100% [1-^13^C_1_]-glucose or 100% [6-^13^C_1_]-glucose as the sole carbon source. The biomass corresponding to 0.5-0.6 mg of CDW was collected in triplicates by fast centrifugation (13,000×*g*, 30 s, −4°C). Bacterial pellets were immediately frozen in liquid N_2_. Samples were then extracted three times with 0.5 mL of 60% (v/v) ethanol buffered with 10 mM ammonium acetate (pH = 7.2) at 78°C for 1 min. After each extraction step, the biomass was separated by centrifugation at 13,000×*g* for 1 min. The three liquid extracts were pooled and dried at 120 μbar, and stored at −80°C thereafter. Samples were re-suspended in 20 μL of MilliQ water, distributed in sealed 96-well microtiter plates, and injected into a Waters Acquity UPLC (Waters Corp., Milford, MA, USA) with a Waters Acquity T3 column (150 mm × 2.1 mm × 1.8 μm, Waters Corp.) coupled to a Thermo TSQ Quantum Ultra triple quadrupole instrument (Thermo Fisher Scientific Inc., Waltham, MA) with electrospray ionization. ^13^C-Labeling patterns of free intracellular metabolites were determined for dihydroxyacetone phosphate (DHAP), F6P, FBP, G6P, 6PG, PEP, R5P, Ru5P, S7P, X5P and Pyr as described previously^69^. All other metabolites are listed in **Supplementary Data 1**. Relative abundance was calculated as the total ion count of all measured ^13^C isotopes for each metabolite. The raw data of labeling experiments is available in **Supplementary Data 1**.

### Determination of ^13^C-labeling patterns by GC-MS

Cultures were grown on either 100% [1-^13^C_1_]-glucose or a mixture of 20% (w/w) [U-^13^C_6_]-glucose and 80% (w/w) naturally-labelled glucose, and 5 mL aliquots of cell broth were harvested by centrifugation at 1,200×*g* and −4°C for 10 min. Bacterial pellets were washed twice with 1 mL of 9 g L^−1^ NaCl, hydrolyzed in 1 mL of 6 M HCl for 24 h at 110°C, and desiccated overnight at 85°C under a constant air stream. The hydrolyzate was dissolved in 50 μL of 99.8% (w/v) dimethyl formamide and subsequently transferred into a new tube. For sample derivatization, 30 μL of *N*-methyl-*N*-(*tert*-butyldimethylsilyl)-trifluoroacetamide was added to the biomass hydrolyzate and incubated at 85°C for 60 min. The ^13^C-labeling patterns of proteinogenic amino acids were determined on a 6890N Network GC system with a 5975 inert XL mass selective detector (Agilent Technologies Inc., Santa Clara, CA, USA) as described previously^70^. The raw GC-MS data from four independent experiments is presented in **Supplementary Data 2**.

### Metabolic flux ratio analysis

Mass distribution vectors of the proteinogenic amino acids were corrected for the natural abundance of all stable isotopes, and the relative metabolic flux ratios [i] oxaloacetate (OAA) from Pyr, [ii] glyoxylate shunt, [iii] PEP from OAA, and [iv] the lower and upper bound for Pyr from malate were calculated using the Fiat Flux software^71^. The mass distribution vectors of the free intracellular metabolites were corrected for the natural abundance of all stable isotopes using MatLab (The Mathworks Inc., Natick, MA, USA). Novel relative flux ratios were defined and calculated as described previously^33^. The fraction of G6P originating from glucose was estimated using data from the experiments using 100% [1-^13^C_1_]-glucose, whereas 6PG from G6P was calculated using data from either 100% [1-^13^C_1_]-glucose or 100% [6-^13^C_1_]-glucose experiments. Formation of F6P through the PP pathway was estimated using data from 100% [6-^13^C_1_]-glucose experiments, and Pyr through the ED pathway was assessed with data from 100% [1-^13^C_1_]-glucose experiments. The metabolic model used for net-flux analysis was based on a master reaction network with 45 reactions and 33 metabolites. Fluxes were calculated using [i] the stoichiometric reaction matrix, [ii] constraints accounting for the ratios from FiatFlux analysis and additionally for the ratios in the initial steps of glucose catabolism as described above, [iii] physiological data, and [iv] precursor requirements for biomass. The experimentally-determined relative flux ratios were translated into constraints as follows (the reaction numbers, *v*x, are defined in **Fig. S1** in the Supplementary Information). The fraction of G6P originating from glucose (*a*) was estimated as *a* = *v*_3_/(*v*_3_ + *v*_16_); the fraction of F6P originating through the PP pathway (*b*) was calculated as *b* = (*v*_14_ + *v*_15_)/(*v*_14_ + *v*_15_ + *v*_17_); and the fraction of 6PG originating from G6P (*c*) was determined as *c* = *v*_7_/(*v*_4_ + *v*_6_ + *v*_7_). For this ratio, the values from the experiments using data from either 100% [1-^13^C_1_]-glucose or [6-^13^C_1_]-glucose were averaged. The fraction of Pyr originating through the ED pathway (*d*) was determined as *d* = *v*_9_/(*v*_9_ + *v*_22_ + *v*_33_ + *v*_34_). The upper and lower bounds of Pyr originating from malate (*e* and *f*, respectively) were obtained according to *e* ≥ *v*_34_/(*v*_9_ + *v*_22_ + *v*_33_ + *v*_34_) and *f* ≤ *v*_34_/(*v*_9_ + *v*_22_ + *v*_33_ + *v*_34_). The fraction of OAA originating from Pyr (*g*) was obtained following *g* = *v*_32_/(*v*_30_ + *v*_32_). Finally, the fraction of Pyr originating from OAA (*h*) was derived from *h* = *v*_33_/(*v*_9_ + *v*_22_ + *v*_33_ + *v*_34_). The determined linear system of mass balances, flux ratios, quantitative physiology data, and biomass requirements was then solved with the *fmincon* function from MatLab using the Netto module from FiatFlux to obtain the net metabolic fluxes as described previously^71^. The values for all the fluxes within the network are provided in **Table S3** in the Supplementary Information.

### Preparation of cell-free extracts and *in vitro* enzymatic assays

Cell-free extracts were prepared from cells harvested by centrifugation from an appropriate culture volume at 4,000×*g* at 4°C for 10 min. Pellets were suspended in 1 volume of 10 mM phosphate-buffered saline (PBS, pH = 7.5, and previously refrigerated) containing 10 mM 2-mercaptoethanol and centrifuged again. Cells were finally resuspended in 0.3-0.5 volume of the same buffer and sonicated intermittently for 6 min in an ice bath. Sonicated cells were centrifuged at 7,500×*g* at 4°C for 30 min, and the total protein concentration in cell extracts was measured by the Bradford method. The activities of Edd, Eda, Glk, Gcd, and Gad were assayed using standard protocols^72,73^. The activity of Zwf and Gnd were assayed under both saturating and non-saturating, *quasi in vivo* conditions using concentrations reflecting intracellular experimentally determined abundance of cofactors and substrates. For those reactions in which more than one enzyme catalyzes the corresponding transformation (e.g., Zwf, for which there are three isozymes in *P. putida* KT2440), the total activity is reported. In the latter case, the concentrations of the substrates (experimentally determined in cell-free extracts of glucose-grown KT2440 by means of LC-MS/MS), were 1.2 mM G6P and 2.4 mM 6PG.

### Glutathione quantification

Glutathione (both oxidized and reduced forms) was quantified in cells harvested from 25 mL-culture samples using a modification of the procedure described by Michie *et al.*^74^. Cells were pelleted by centrifugation (4,000×*g* at 4°C for 10 min) and suspended in 2 mL of cold PBS. The cell suspension was mixed 1:1 (v/v) with a freshly-prepared 10% (w/v) 5-sulfosalicylic acid solution, incubated on an ice bath for 10 min and then sonicated. For the measurement of total glutathione, 40 μL of Tris(2-carboxyethyl)phosphine·HCl was added to an equal volume of the cell-free extract. For the measurement of reduced glutathione, 40 μL of H_2_O were used instead. Samples were centrifuged at 15,600×*g* for 5 min after treatment, and 25 μL of the supernatant fluid was added to 100 μL of a buffer containing 200 mM *N*-ethylmorpholine and 20 mM NaOH. Following the addition of 50 μL of 0.5 N NaOH to this mixture, glutathione was derivatized by addition of 10 μL of 10 mM naphthalene-2,3-dicarboxaldehyde. The mixture was incubated at room temperature for 30 min, and the fluorescence intensity of the resulting naphthalene-2,3-dicarboxaldehyde–glutathione conjugate was measured at an excitation wavelength (*λ*_excitation_) = 472 nm and an emission wavelength (*λ*_excitation_) = 528 nm. A standard curve was constructed with known amounts of freshly-prepared naphthalene-2,3-dicarboxaldehyde–glutathione conjugate.

### Chemicals and enzymes

[1-^13^C_1_]-Glucose and [6-^13^C_1_]-glucose were purchased from Cambridge Isotope Laboratories Inc. (Tewksbury, MA, USA), and [U-^13^C_6_]-glucose was purchased from Sigma-Aldrich Co. (St. Louis, MO, USA). Other chemicals and enzymes used for *in vitro* assays were obtained from Sigma-Aldrich Co. and Merck KGaA (Darmstadt, Germany). H_2_O_2_ solutions were freshly prepared from a 30% (w/w) stock solution (Sigma-Aldrich Co.).

### Data availability

Metabolite levels and ^13^C-labelling data are available in the Supplementary Information files. Other datasets generated and analyzed in this study are available from the corresponding authors on request.

## Acknowledgements

This work was funded by The Novo Nordisk Foundation (individual grant NNF10CC1016517, and *LiFe*, NNF18OC0034818), the European Union’s *Horizon 2020* Research and Innovation Programme under grant agreement No. 814418 (*SinFonia*) and the Danish Council for Independent Research (*SWEET*, DFF-Research Project 8021-00039B) to P.I.N. This work was also funded by the *MADONNA* (H2020-FET-OPEN-RIA-2017-1-766975), *BioRoboost* (H2020-NMBP-BIO-CSA-2018), *SYNBIO4FLAV* (H2020-NMBP/0500) and *MIX-UP* (H2020-Grant 870294) Contracts of the European Union and the S2017/BMD-3691 *InGEMICS-CM* Project of the Comunidad Autónoma de Madrid (European Structural and Investment Funds) to V.D.L.

## Author contributions

P.I.N. and V.D.L. conceived the study and wrote the manuscript with input from all other authors. P.I.N., T.F., M.C. and A.S.P. carried out the quantitative physiology and tracer experiments. P.I.N. and A.S.P. measured the in vitro enzymatic activities. T.F. performed metabolomics and fluxome analysis. All authors contributed to the discussion of the experimental results.

## Additional information

Supplementary Information accompanies this paper online.

## References

1. Imlay, J.A. Where in the world do bacteria experience oxidative stress? Environ. Microbiol. 21, 521–530 (2019).

2. Finkel, T. Oxidant signals and oxidative stress. Curr. Opin. Cell Biol. 15, 247–254 (2003).

3. Mishra, S. & Imlay, J. Why do bacteria use so many enzymes to scavenge hydrogen peroxide? Arch. Biochem. Biophys. 525, 145–160 (2012).

4. Chaput, V., Martin, A. & Lejay, L. Redox metabolism: the hidden player in carbon and nitrogen signaling? J. Exp. Bot., In press, DOI: 10.1093/jxb/eraa1078 (2020).

5. Cadet, J., Douki, T., Gasparutto, D. & Ravanat, J.L. Oxidative damage to DNA: formation, measurement and biochemical features. Mut. Res. 531, 5–23 (2003).

6. Imlay, J.A., Sethu, R. & Rohaun, S.K. Evolutionary adaptations that enable enzymes to tolerate oxidative stress. Free Radic. Biol. Med. 140, 4–13 (2019).

7. Arts, I.S., Gennaris, A. & Collet, J.F. Reducing systems protecting the bacterial cell envelope from oxidative damage. FEBS Lett. 589, 1559–1568 (2015).

8. Dan Dunn, J., Álvarez, L.A.J., Zhang, X. & Soldati, T. Reactive oxygen species and mitochondria: A nexus of cellular homeostasis. Redox Biol. 6, 472–485 (2015).

9. Imlay, J.A. The molecular mechanisms and physiological consequences of oxidative stress: lessons from a model bacterium. Nat. Rev. Microbiol. 11, 443–454 (2013).

10. Jamieson, D.J. The effect of oxidative stress on *Saccharomyces cerevisiae*. Redox Rep. 1, 89–95 (1995).

11. Eleutherio, E. et al. Oxidative stress and aging: Learning from yeast lessons. Fungal Biol. 122, 514–525 (2018).

12. Brown, A.J.P., Cowen, L.E., di Pietro, A. & Quinn, J. Stress adaptation. Microbiol. Spectr. 5 (2017).

13. Zheng, M. et al. DNA microarray-mediated transcriptional profiling of the *Escherichia coli* response to hydrogen peroxide. J. Bacteriol. 183, 4562–4570 (2001).

14. Gorrini, C., Harris, I.S. & Mak, T.W. Modulation of oxidative stress as an anticancer strategy. Nat. Rev. Drug Discov. 12, 931–947 (2013).

15. Zhang, J. et al. ROS and ROS-mediated cellular signaling. Oxid. Med. Cell Longev. 2016, 4350965 (2016).

16. Balsa, E. et al. Defective NADPH production in mitochondrial disease complex I causes inflammation and cell death. Nat. Commun. 11, 2714 (2020).

17. Boronat, S. et al. Thiol-based H_2_O_2_ signalling in microbial systems. Redox Biol. 2, 395–399 (2014).

18. Doroshow, J.H. Glutathione peroxidase and oxidative stress. Toxicol. Lett. 82–83, 395–398 (1995).

19. Chechik, G. et al. Activity motifs reveal principles of timing in transcriptional control of the yeast metabolic network. Nat. Biotechnol. 26, 1251–1259 (2008).

20. Fuhrer, T. & Sauer, U. Different biochemical mechanisms ensure network-wide balancing of reducing equivalents in microbial metabolism. J. Bacteriol. 191, 2112–2121 (2009).

21. Rui, B. et al. A systematic investigation of *Escherichia coli* central carbon metabolism in response to superoxide stress. BMC Syst. Biol. 4, 122 (2010).

22. Kuehne, A. et al. Acute activation of oxidative pentose phosphate pathway as first-line response to oxidative stress in human skin cells. Mol. Cell 59, 359–371 (2015).

23. Christodoulou, D. et al. Reserve flux capacity in the pentose phosphate pathway enables *Escherichia coli*’s rapid response to oxidative stress. Cell Syst. 6, 569–578 (2018).

24. Belda, E. et al. The revisited genome of *Pseudomonas putida* KT2440 enlightens its value as a robust metabolic *chassis*. Environ. Microbiol. 18, 3403–3424 (2016).

25. Nogales, J. et al. High-quality genome-scale metabolic modelling of *Pseudomonas putida* highlights its broad metabolic capabilities. Environ. Microbiol. 22, 255–269 (2020).

26. Nelson, K.E. et al. Complete genome sequence and comparative analysis of the metabolically versatile *Pseudomonas putida* KT2440. Environ. Microbiol. 4, 799–808 (2002).

27. Volke, D.C., Calero, P. & Nikel, P.I. Pseudomonas putida. Trends Microbiol. 28, 512–513 (2020).

28. Nikel, P.I., Chavarría, M., Danchin, A. & de Lorenzo, V. From dirt to industrial applications: *Pseudomonas putida* as a Synthetic Biology *chassis* for hosting harsh biochemical reactions. Curr. Opin. Chem. Biol. 34, 20–29 (2016).

29. Akkaya, Ö., Pérez-Pantoja, D., Calles, B., Nikel, P.I. & de Lorenzo, V. The metabolic redox regime of *Pseudomonas putida* tunes its evolvability toward novel xenobiotic substrates. mBio 9, e01512–01518 (2018).

30. Nikel, P.I., Martínez-García, E. & de Lorenzo, V. Biotechnological domestication of pseudomonads using synthetic biology. Nat. Rev. Microbiol. 12, 368–379 (2014).

31. del Castillo, T. et al. Convergent peripheral pathways catalyze initial glucose catabolism in *Pseudomonas putida*: genomic and flux analysis. J Bacteriol 189, 5142–5152 (2007).

32. Sudarsan, S., Dethlefsen, S., Blank, L.M., Siemann-Herzberg, M. & Schmid, A. The functional structure of central carbon metabolism in *Pseudomonas putida* KT2440. Appl. Environ. Microbiol. 80, 5292–5303 (2014).

33. Nikel, P.I., Chavarría, M., Fuhrer, T., Sauer, U. & de Lorenzo, V. *Pseudomonas putida* KT2440 strain metabolizes glucose through a cycle formed by enzymes of the Entner-Doudoroff, Embden-Meyerhof-Parnas, and pentose phosphate pathways. J. Biol. Chem. 290, 25920–25932 (2015).

34. Kohlstedt, M. & Wittmann, C. GC-MS-based ^13^C metabolic flux analysis resolves the parallel and cyclic glucose metabolism of *Pseudomonas putida* KT2440 and *Pseudomonas aeruginosa* PAO1. Metab. Eng. 54, 35–53 (2019).

35. Kukurugya, M.A. et al. Multi-omics analysis unravels a segregated metabolic flux network that tunes co-utilization of sugar and aromatic carbons in *Pseudomonas putida*. J. Biol. Chem. 294, 8464–8479 (2019).

36. Cabiscol, E., Tamarit, J. & Ros, J. Oxidative stress in bacteria and protein damage by reactive oxygen species. Internatl. Microbiol. 3, 3–8 (2000).

37. Fuhrer, T., Fischer, E. & Sauer, U. Experimental identification and quantification of glucose metabolism in seven bacterial species. J. Bacteriol. 187, 1581–1590 (2005).

38. Chavarría, M., Nikel, P.I., Pérez-Pantoja, D. & de Lorenzo, V. The Entner-Doudoroff pathway empowers *Pseudomonas putida* KT2440 with a high tolerance to oxidative stress. Environ. Microbiol. 15, 1772–1785 (2013).

39. Sánchez-Pascuala, A., de Lorenzo, V. & Nikel, P.I. Refactoring the Embden-Meyerhof-Parnas pathway as a whole of portable *GlucoBricks* for implantation of glycolytic modules in Gram-negative bacteria. ACS Synth. Biol. 6, 793–805 (2017).

40. Mailloux, R.J., Lemire, J. & Appanna, V.D. Metabolic networks to combat oxidative stress in *Pseudomonas fluorescens*. Antonie van Leeuwenhoek 99, 433–442 (2011).

41. Calero, P. & Nikel, P.I. Chasing bacterial *chassis* for metabolic engineering: A perspective review from classical to non-traditional microorganisms. Microb. Biotechnol. 12, 98–124 (2019).

42. Shimizu, K. & Matsuoka, Y. Redox rebalance against genetic perturbations and modulation of central carbon metabolism by the oxidative stress regulation. Biotechnol. Adv. 37, 107441 (2019).

43. Chakraborty, S. et al. Glycolytic reprograming in *Salmonella* counters NOX2-mediated dissipation of ∆pH. Nat. Commun. 11, 1783–1783 (2020).

44. Gottesman, S. Trouble is coming: Signaling pathways that regulate general stress responses in bacteria. J. Biol. Chem. 294, 11685–11700 (2019).

45. Kim, J. & Park, W. Oxidative stress response in *Pseudomonas putida*. Appl. Microbiol. Biotechnol. 98, 6933–6946 (2014).

46. MacLean, A., Bley, A.M., Appanna, V.P. & Appanna, V.D. Metabolic manipulation by *Pseudomonas fluorescens*: a powerful stratagem against oxidative and metal stress. J. Med. Microbiol. 69, 339–346 (2020).

47. Bojanovič, K., D’Arrigo, I. & Long, K.S. Global transcriptional responses to osmotic, oxidative, and imipenem stress conditions in *Pseudomonas putida*. Appl. Environ. Microbiol. 83, e03236–03216 (2017).

48. Ralser, M. et al. Dynamic rerouting of the carbohydrate flux is key to counteracting oxidative stress. J. Biol. 6, 10 (2007).

49. Stincone, A. et al. The return of metabolism: biochemistry and physiology of the pentose phosphate pathway. Biol. Rev. 90, 927–963 (2015).

50. Christodoulou, D. et al. Reserve flux capacity in the pentose phosphate pathway by NADPH binding is conserved across kingdoms. iScience 19, 1133–1144 (2019).

51. Chavarría, M., Kleijn, R.J., Sauer, U., Pflüger-Grau, K. & de Lorenzo, V. Regulatory tasks of the phosphoenolpyruvate-phosphotransferase system of *Pseudomonas putida* in central carbon metabolism. mBio 3, e00028–00012 (2012).

52. Zhou, H. et al. Modulation of *Pseudomonas aeruginosa* quorum sensing by glutathione. J. Bacteriol. 201, e00685–00618 (2019).

53. Wongsaroj, L. et al. *Pseudomonas aeruginosa* glutathione biosynthesis genes play multiple roles in stress protection, bacterial virulence and biofilm formation. PLoS One 13, e0205815 (2018).

54. Lemire, J., Alhasawi, A., Appanna, V.P., Tharmalingam, S. & Appanna, V.D. Metabolic defence against oxidative stress: The road less travelled so far. J. Appl. Microbiol. 123, 798–809 (2017).

55. Staerck, C. et al. Microbial antioxidant defense enzymes. Microb. Pathog. 110, 56–65 (2017).

56. Hishinuma, S., Yuki, M., Fujimura, M. & Fukumori, F. OxyR regulated the expression of two major catalases, KatA and KatB, along with peroxiredoxin, AhpC in *Pseudomonas putida*. Environ. Microbiol. 8, 2115–2124 (2006).

57. Heim, S. et al. Proteome reference map of *Pseudomonas putida* strain KT2440 for genome expression profiling: distinct responses of KT2440 and *Pseudomonas aeruginosa* strain PAO1 to iron deprivation and a new form of superoxide dismutase. Environ. Microbiol. 5, 1257–1269 (2003).

58. Yeom, J., Jeon, C.O., Madsen, E.L. & Park, W. *In vitro* and *in vivo* interactions of ferredoxin-NADP^+^ reductases in *Pseudomonas putida*. J. Biochem. 145, 481–491 (2009).

59. Nikel, P.I., Chavarría, M., Martínez-García, E., Taylor, A.C. & de Lorenzo, V. Accumulation of inorganic polyphosphate enables stress endurance and catalytic vigour in *Pseudomonas putida* KT2440. Microb. Cell Fact. 12, 50 (2013).

60. Nikel, P.I., Pérez-Pantoja, D. & de Lorenzo, V. Pyridine nucleotide transhydrogenases enable redox balance of *Pseudomonas putida* during biodegradation of aromatic compounds. Environ. Microbiol. 18, 3565–3582 (2016).

61. Ralser, M. et al. Metabolic reconfiguration precedes transcriptional regulation in the antioxidant response. Nat. Biotechnol. 27, 604–605 (2009).

62. Akkaya, Ö., Nikel, P.I., Pérez-Pantoja, D. & de Lorenzo, V. Evolving metabolism of 2,4-dinitrotoluene triggers SOS-independent diversification of host cells. Environ. Microbiol. 21, 314–326 (2019).

63. Pérez-Pantoja, D., Nikel, P.I., Chavarría, M. & de Lorenzo, V. Endogenous stress caused by faulty oxidation reactions fosters evolution of 2,4-dinitrotoluene-degrading bacteria. PLoS Genet. 9, e1003764 (2013).

64. Ankenbauer, A. et al. *Pseudomonas putida* KT2440 is naturally endowed to withstand industrial-scale stress conditions. Microb. Biotechnol. 13, 1145–1161 (2020).

65. Bagdasarian, M. et al. Specific purpose plasmid cloning vectors. II. Broad host range, high copy number, RSF1010-derived vectors, and a host-vector system for gene cloning in *Pseudomonas*. Gene 16, 237–247 (1981).

66. Sambrook, J. & Russell, D.W. Molecular cloning: a laboratory manual, Edn. 3rd. (Cold Spring Harbor Laboratory, Cold Spring Harbor; 2001).

67. Nikel, P.I., Pettinari, M.J., Ramírez, M.C., Galvagno, M.A. & Méndez, B.S. *Escherichia coli arcA* mutants: metabolic profile characterization of microaerobic cultures using glycerol as a carbon source. J. Mol. Microbiol. Biotechnol. 15, 48–54 (2008).

68. Nikel, P.I. & Chavarría, M. in Hydrocarbon and Lipid Microbiology Protocols–Synthetic and Systems Biology - Tools. (eds. T.J. McGenity, K.N. Timmis & B. Nogales-Fernández) 39–70 (Humana Press, Heidelberg, Germany; 2016).

69. Rühl, M. et al. Collisional fragmentation of central carbon metabolites in LC-MS/MS increases precision of ¹³C metabolic flux analysis. Biotechnol. Bioeng. 109, 763–771 (2012).

70. Nanchen, A., Fuhrer, T. & Sauer, U. Determination of metabolic flux ratios from ^13^C-experiments and gas chromatography-mass spectrometry data: protocol and principles. Methods Mol. Biol. 358, 177–197 (2007).

71. Zamboni, N., Fischer, E. & Sauer, U. FiatFlux – a software for metabolic flux analysis from ^13^C-glucose experiments. BMC Bioinformatics 6, 209 (2005).

72. Sánchez-Pascuala, A., Fernández-Cabezón, L., de Lorenzo, V. & Nikel, P.I. Functional implementation of a linear glycolysis for sugar catabolism in *Pseudomonas putida*. Metab. Eng. 54, 200–211 (2019).

73. Corona, F., Martínez, J.L. & Nikel, P.I. The global regulator Crc orchestrates the metabolic robustness underlying oxidative stress resistance in *Pseudomonas aeruginosa*. Environ. Microbiol. 21, 898–912 (2019).

74. Michie, K.L. et al. The role of *Pseudomonas aeruginosa* glutathione biosynthesis in lung and soft tissue infection. Infect. Immun., In press, DOI: 10.1128/iai.00116-00120 (2020).

