## Supplementary Information for "Redox stress reshapes carbon fluxes of *Pseudomonas putida* for cytosolic glucose oxidation and NADPH generation"

### Redox stress reshapes carbon fluxes of the core metabolic cycle of *Pseudomonas putida* for adaptation to oxidative environments

Pablo I. Nikel, Tobias Fuhrer, Max Chavarría, Alberto Sánchez-Pascuala, Uwe Sauer, and Víctor de Lorenzo

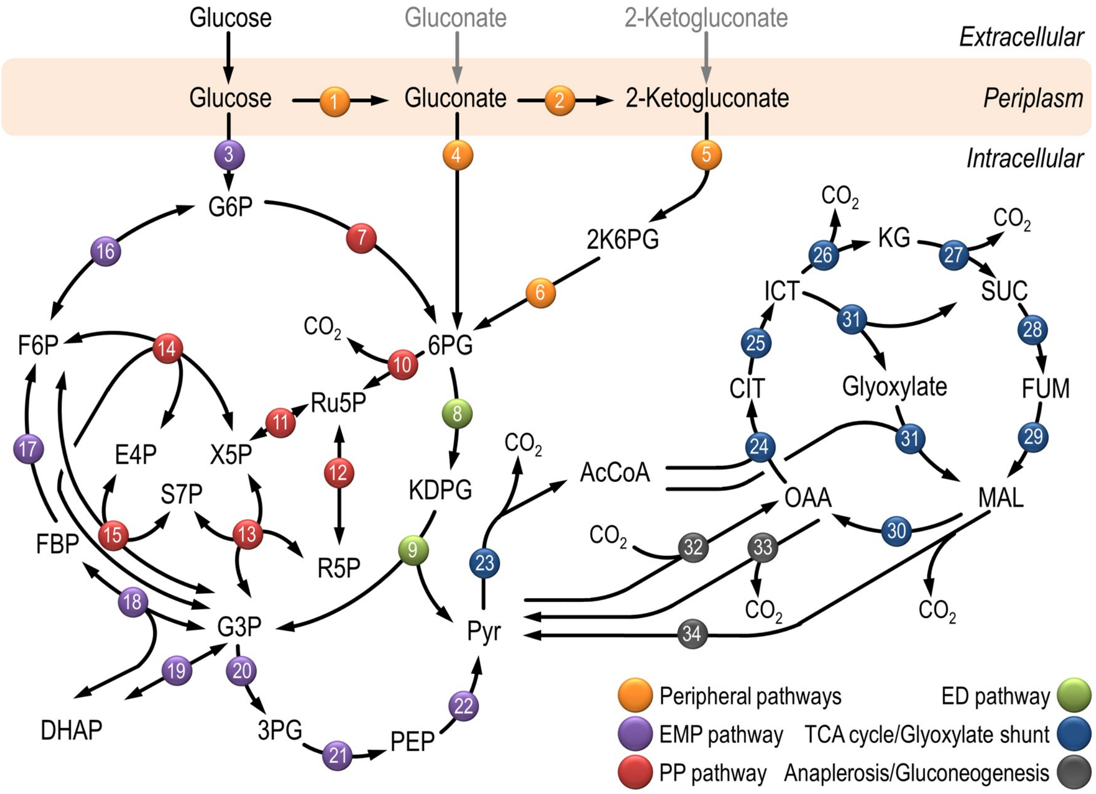


**Figure S1. Biochemical pathways involved in glucose catabolism in *Pseudomonas putida* KT2440.** The metabolic network is sketched around six main metabolic blocks, identified with different colors: [i] peripheral pathways, i.e. oxidative transformation of glucose into gluconate and 2-ketogluconate (and their corresponding phosphorylated derivatives); [ii] Embden-Meyerhof-Parnas (EMP) pathway (non-functional, since 6-phosphofructo-1-kinase is absent); [iii] pentose phosphate (PP) pathway; [iv] Entner-Doudoroff (ED) pathway; [v] tricarboxylic acid (TCA) cycle and glyoxylate shunt; and [vi] anaplerotic and gluconeogenic reactions. The oxidative transformations in the outer membrane and in the periplasmic space are shown at the top of the scheme, along with the transport systems for glucose, gluconate and 2-ketogluconate into the cytoplasm. Some reactions have been lumped to simplify the diagram. Transport of gluconate and 2-ketogluconate from the extracellular space is indicated by gray arrows. All the reactions in the network have been assigned a number (circled), and the complete list of the enzymes and isozymes catalyzing each step is listed in **Table S1**. Abbreviations are as follows: G6P, glucose-6-phosphate; F6P, fructose-6-phosphate; FBP, fructose-1,6-bisphosphate; DHAP, dihydroxyacetone phosphate; 6PG, 6-phosphogluconate; KDPG, 2-keto-3-deoxy-6-phosphogluconate; 2K6PG, 2-keto-6-phosphogluconate; Ru5P, ribulose-5-phosphate; R5P, ribose-5-phosphate; X5P, xylulose-5-phosphate; S7P, sedoheptulose-7-phosphate; E4P, erythrose-4-phosphate; G3P, glyceraldehyde-3-phosphate; 3PG, 3-phosphoglycerate; PEP, phosphoenolpyruvate; Pyr, pyruvate; AcCoA, acetyl-coenzyme A; OAA, oxaloacetate; CIT, citrate; ICT, isocitrate; KG, 2-ketoglutarate; SUC, succinate; FUM, fumarate; and MAL, malate.

**Table S1.** Components of the biochemical network of *Pseudomonas putida* KT2440[a].

| Block | Code | Reaction | Enzyme(s) | Name(s) and PP number(s) |
|---|---|---|---|---|
| Peripheral pathways | 1 | Glucose + Ubiquinone ⟶ Glucono-1,5-lactone + Ubiquinol | Glucose dehydrogenase | Gcd (PP_1444) |
| | | Glucono-1,5-lactone + $H_2O$ ⟶ Gluconate + $H^+$ | Gluconolactonase | Gnl (PP_1170) |
| | 2 | Gluconate + Ubiquinone ⟶ 2-Ketogluconate + Ubiquinol | Gluconate 2-dehydrogenase | PP_3382 PP_3383 PP_3384 |
| | 4 | Gluconate + ATP ⟶ 6PG + ADP + $H^+$ | Gluconate kinase | GnuK (PP_3416) |
| | 5 | 2-Ketogluconate + ATP ⟶ 2K6PG + ADP + $H^+$ | 2-Ketoglucokinase | KguK (PP_3378) |
| | 6 | 2K6PG + NADPH + $H^+$ ⟶ 6PG + $NADP^+$ | 2-Ketogluconate-6-phosphate reductase | KguD (PP_3376) |
| Pentose phosphate pathway | 7 | G6P + $NADP^+$ ⟶ 6-Phosphoglucono-1,5-lactone + NADPH + $H^+$ | Glucose-6-phosphate 1-dehydrogenase | ZwfA (PP_1022) ZwfB (PP_4042) Zwf (PP_5351) |
| | | 6-Phosphoglucono-1,5-lactone + $H_2O$ ⟶ 6PG + $H^+$ | 6-Phospho-gluconolactonase | Pgl (PP_1023) |
| | 10 | 6PG + $NADP^+$ ⟶ Ru5P + NADPH + $CO_2$ | 6-Phosphogluconate dehydrogenase | GntZ (PP_4043) |
| | 11 | Ru5P ↔ Xu5P | Ribulose-5-phosphate 3-epimerase | Rpe (PP_0415) |
| | 12 | Ru5P ↔ Ri5P | Ribose-5-phosphate isomerase | RpiA (PP_5150) |
| | 13 | Xu5P + R5P ↔ S7P + G3P | Transketolase | TktA (PP_4965) |
| | 14 | Xu5P + E4P ↔ G3P + F6P | | |
| | 15 | S7P + G3P ↔ E4P + F6P | Transaldolase | Tal (PP_2168) |
| Entner-Doudoroff pathway | 8 | 6PG ⟶ KDPG + $H_2O$ | 6-Phosphogluconate dehydratase | Edd (PP_1010) |
| | 9 | KDPG ⟶ G3P + Pyr | 2-Keto-3-deoxy-6-phosphogluconate aldolase | Eda (PP_1024) |

| | | | | |
|---|---|---|---|---|
| Embden-Meyerhof-Parnas pathway | 3 | Glucose + ATP $\rightarrow$ G6P + ADP + $H^+$ | Glucokinase | Glk (PP_1011) |
| | 16 | G6P $\rightarrow$ F6P | Glucose-6-phosphate isomerase | Pgi-I (PP_1808) Pgi-II (PP_4701) |
| | 17 | FBP + $H_2O$ + ADP $\rightarrow$ F6P + ATP | Fructose-1,6-bisphosphatase | Fbp (PP_5040) |
| | 18 | DHAP + G3P $\leftrightarrow$ FBP | Fructose-1,6-bisphosphate aldolase | Fda (PP_4960) PP_2871 PP_3224 |
| | 19 | G3P $\leftrightarrow$ DHAP | Triose phosphate isomerase | TpiA (PP_4715) |
| | 20 | G3P + $NAD^+$ + Pi $\rightarrow$ 1,3-Bisphosphoglycerate + NADH + $H^+$ | Glyceraldehyde-3-phosphate dehydrogenase | GapA (PP_1009) GapB (PP_2149) PP_0665 PP_3443 |
| | | 1,3-Bisphosphoglycerate + ADP $\rightarrow$ 3PG + ATP | Phosphoglycerate kinase | Pgk (PP_4963) |
| | 21 | 3PG $\rightarrow$ 2PG | Phosphoglycerate mutase | GpmI (PP_5056) PP_2243 PP_3923 PP_4450 |
| | | 2PG $\rightarrow$ PEP + $H_2O$ | Enolase | Eno (PP_1612) |
| | 22 | PEP + ADP + $H^+$ $\rightarrow$ Pyr + ATP | Pyruvate kinase | PykA (PP_1362) Pyk (PP_4301) |
| Tricarboxylic acid cycle / Glyoxylate shunt | 23 | Pyr + $NAD^+$ + Coenzyme A $\rightarrow$ AcCoA + NADH + $CO_2$ | Pyruvate dehydrogenase | AcoA (PP_0555) AcoC (PP_0553) AceF (PP_0338) AceE (PP_0339) |
| | 24 | OAA + AcCoA + $H_2O$ $\rightarrow$ CIT + Coenzyme A + $H^+$ | Citrate synthase | GltA (PP_4194) |
| | 25 | CIT $\rightarrow$ ICT | Aconitate hydratase | AcnA-I (PP_2112) AcnA-II (PP_2336) AcnB (PP_2339) |

| | | | | |
|---|---|---|---|---|
| Tricarboxylic acid cycle / Glyoxylate shunt | 26 | ICT + $NADP^+$ ⟶ KG + $CO_2$ + NADPH + $H^+$ | Isocitrate dehydrogenase | Icd (PP_4011) Idh (PP_4012) |
| | 27 | KG + Coenzyme A + $NAD^+$ ⟶ Succinyl-Coenzyme A + NADH + $H^+$ + $CO_2$ | 2-Ketoglutarate dehydrogenase | Lpd (PP_5366) LpdG (PP_4187) LpdV (PP_4404) SucA (PP_4189) SucB (PP_4188) PP2652 PP3662 |
| | | Succinyl-Coenzyme A + ADP + Pi ⟶ SUC + Coenzyme A + ATP | Succinyl-coenzyme A synthetase | SucC (PP_4186) SucD (PP_4185) ScpC (PP_0154) |
| | 28 | SUC + Ubiquinone ⟶ FUM + Ubiquinol | Succinate dehydrogenase | SdhA (PP_4191) SdhB (PP_4190) SdhC (PP_4193) SdhD (PP_4192) |
| | 29 | FUM + $H_2O$ ⟶ MAL | Fumarate hydratase | FumC-I (PP_0944) FumC-II (PP_1755) PP_0897 |
| | 30 | MAL + $NAD^+$ (quinone) ⟶ OAA + NADH (quinol) + $H^+$ | Malate dehydrogenase / Malate:quinone oxidoreductase | Mdh (PP_0654) PP_3591 Mqo-1 (PP_0751) Mqo-2 (PP_1251) Mqo-3 (PP2925) |
| | 31 | ICT ⟶ SUC + Glyoxylate | Isocitrate lyase | AceA (PP_4116) |
| | | Glyoxylate + AcCoA + $H_2O$ ⟶ MAL + Coenzyme A + $H^+$ | Malate synthase | GlcB (PP_0356) |

| | | | | |
|---|---|---|---|---|
| Anaplerosis / Gluconeogenesis | 32 | Pyr + $CO_2$ ⟶ OAA + $H^+$ | Pyruvate carboxylase | PycA (PP_5347) PycB (PP_5346) |
| | 33 | OAA + Pi ⟶ PEP + $CO_2$ | Phosphoenol-pyruvate carboxylase[d] | Ppc (PP_1505) |
| | 34 | MAL + $NADP^+$ ⟶ Pyr + $CO_2$ + NADPH | Malic enzyme | MaeB (PP_5085) |

[a] Information compiled from the *Pseudomonas* Genome Database[1,2], MetaCyc[3] and the literature[4-6]. In the instances in which no gene name has been assigned, the PP number is given for each open reading frame. Biochemical reactions are coded according to the six functional blocks indicated in **Fig. S1**. According to Nelson *et al*.[7], *pckA* (*PP_0253*, encoding phosphoenolpyruvate carboxykinase) contains an authentic frameshift and therefore the open reading frame is classified as a pseudogene in the *Pseudomonas* Genome Database. All abbreviations are defined in the legend to **Fig. S1**. Pi, inorganic orthophosphate.

**Table S2.** Selected metabolic flux ratios used for ratio-constrained flux balance analysis[a].

| Metabolic flux ratio | Code(s) in **Fig. S1** | Ratio (mean ± SD)[e] | |
|---|---|---|---|
| | | Control conditions | + $H_2O_2$ |
| G6P from glucose[b] | 3 | 0.53 ± 0.06 | 0.43 ± 0.01 |
| 6PG from G6P[b] | 7 | 0.14 ± 0.02 | 0.56 ± 0.01 |
| 6PG from G6P[c] | 7 | 0.17 ± 0.02 | 0.56 ± 0.05 |
| F6P from the PP pathway[c] | 14, 15 | 0.23 ± 0.03 | 0.79 ± 0.02 |
| Pyruvate through the ED pathway[b] | 9 | 0.44 ± 0.02 | 0.51 ± 0.03 |
| Glyoxylate shunt[d] | 31 | N.D. | N.D. |
| OAA from pyuvate[d] | 32 | 0.65 ± 0.06 | 0.63 ± 0.06 |
| Phosphoenolpyruvate from oxaloacetate[d] | 33 | 0.00 ± 0.04 | 0.01 ± 0.04 |
| Pyruvate from malate (UB)[d] | 34 | 0.71 ± 0.16 | 0.65 ± 0.14 |
| Pyruvate from malate (LB)[d] | 34 | 0.25 ± 0.04 | 0.24 ± 0.03 |

[a] Flux ratios are shaded according to the metabolic block they belong according to **Fig. S1** and **Table S1**. Abbreviations: G6P, glucose-6-phosphate; 6PG, 6-phosphogluconate; F6P, fructose-6-phosphate; PP pathway, pentose phosphate pathway; ED pathway, Entner-Doudoroff pathway; UB, upper bound; LB, lower bound; SD, standard deviation; and ND, not detected.

[b] Determined from 100% [1-$^{13}$C]-glucose experiments.

[c] Determined from 100% [6-$^{13}$C]-glucose experiments.

[d] Determined from 20% [U-$^{13}C_6$]-glucose experiments.

[e] Standard deviations (SD) for each relative metabolic flux ratio were calculated using the covariance matrices of the respective mass distribution vectors by applying the Gaussian law of error propagation.

**Table S3.** Net flux values for central metabolic pathways[a] of *P. putida* KT2440 grown on glucose under control and oxidative stress conditions.

| Functional block | Reaction | **Flux value** (mmol $g_{CDW}^{-1}$ $h^{-1}$) **± standard error** | |
|---|---|---|---|
| | | **Control conditions** | **+ $H_2O_2$** |
| Peripheral pathways | 1 | 5.53 ± 0.05 | 3.96 ± 0.07 |
| | 2 | 0.72 ± 0.04 | 0.52 ± 0.04 |
| | 4 | 4.79 ± 0.07 | 3.45 ± 0.07 |
| | 5 | 0.72 ± 0.04 | 0.52 ± 0.04 |
| | 6 | 0.72 ± 0.04 | 0.52 ± 0.04 |
| Pentose phosphate pathway | 7 | 1.17 ± 0.08 | 5.02 ± 0.05 |
| | 10 | 0.59 ± 0.03 | 3.89 ± 0.13 |
| | 11 | 0.15 ± 0.59 | 2.31 ± 3.89 |
| | 12 | 0.44 ± 0.59 | 1.58 ± 3.89 |
| | 13 | 0.15 ± 0.01 | 1.28 ± 0.05 |
| | 14 | 0.01 ± 0.01 | 1.04 ± 0.05 |
| | 15 | 0.15 ± 0.01 | 1.28 ± 0.05 |
| Entner-Doudoroff pathway | 8 | 6.12 ± 0.05 | 5.09 ± 0.05 |
| | 9 | 6.12 ± 0.05 | 5.09 ± 0.05 |
| Embden-Meyerhof-Parnas pathway | 3 | 0.63 ± 0.04 | 2.18 ± 0.07 |
| | 16 | 0.57 ± 0.04 | 2.87 ± 0.11 |
| | 17 | 0.46 ± 0.03 | 0.61 ± 0.04 |
| | 18 | 0.46 ± 0.03 | 0.61 ± 0.04 |
| | 19 | 0.46 ± 0.03 | 0.61 ± 0.04 |
| | 20 | 5.11 ± 0.04 | 4.86 ± 0.05 |
| | 21 | 4.36 ± 0.07 | 4.03 ± 0.07 |
| | 22 | 3.89 ± 0.08 | 3.53 ± 0.08 |
| Tricarboxylic acid cycle / Glyoxylate shunt | 23 | 6.67 ± 0.18 | 5.01 ± 0.17 |
| | 24 | 5.44 ± 0.21 | 3.67 ± 0.18 |
| | 25 | 5.44 ± 0.21 | 3.67 ± 0.18 |
| | 26 | 5.44 ± 0.21 | 3.67 ± 0.18 |
| | 27 | 4.71 ± 0.22 | 2.85 ± 0.21 |
| | 28 | 4.71 ± 0.22 | 2.85 ± 0.21 |
| | 29 | 4.71 ± 0.22 | 2.85 ± 0.21 |
| | 30 | 2.01 ± 0.11 | 1.75 ± 0.09 |
| | 31 | 0.00 ± 0.00 | 0.00 ± 0.00 |
| Anaplerosis / Gluconeogenesis | 32 | 4.44 ± 0.15 | 2.89 ± 0.12 |
| | 33 | 0.15 ± 0.13 | 0.04 ± 0.11 |
| | 34 | 2.69 ± 0.18 | 1.12 ± 0.15 |

[a] The classification and codes of the biochemical reactions is the same as depicted in **Fig. S1** and **Table S1** in the Supplementary Information. The distribution of normalized values for each metabolic flux normalized to the specific rate of glucose consumption is shown in **Fig. 3** in the main text.

**Table S4.** Cofactor specificity for the main dehydrogenases in the central metabolism of *Pseudomonas putida* KT2440.

| Enzyme | Enzyme(s) | Relative cofactor specificity (%) under | | | |
|---|---|---|---|---|---|
| | | Saturating conditions | | Non-saturating, *quasi in vivo* conditions | |
| | | $NAD^+$ | $NADP^+$ | $NAD^+$ | $NADP^+$ |
| G6P dehydrogenase | ZwfA (PP_1022)<br>ZwfB (PP_4042)<br>Zwf (PP_5351) | 32.8 ± 3.6 | 67.2 ± 9.5 | 6.3 ± 0.5 | 93.7 ± 1.1 |
| 6PG dehydrogenase | GntZ (PP_4043) | 23.9 ± 0.8 | 76.1 ± 3.2 | 8.6 ± 0.7 | 91.4 ± 1.2 |
| ICT dehydrogenase | Icd (PP_4011)<br>Idh (PP_4012) | 11.8 ± 1.3 | 88.2 ± 3.3 | 11.1 ± 0.5 | 88.9 ± 2.6 |
| MAL dehydrogenase | Mdh (PP_0654)<br>PP_5391 | 98.4 ± 1.1 | 1.6 ± 0.3 | 97.5 ± 2.9 | 2.5 ± 0.9 |
| 2K6PG reductase | KguD (PP_3376) | 13.3 ± 0.5 | 86.7 ± 1.3 | 10.2 ± 0.8 | 89.8 ± 1.3 |
| G3P dehydrogenase | GapA (PP_1009)<br>GapB (PP_2149)<br>PP_0665<br>PP_3443 | 84.1 ± 7.2 | 15.9 ± 1.3 | 66.8 ± 0.9 | 33.2 ± 0.1 |
| Malic enzyme | MaeB (PP_5085) | 4.6 ± 1.6 | 95.4 ± 2.4 | 3.6 ± 0.1 | 96.4 ± 2.7 |

[a] Values represent the mean of the relative cofactor specificity ± standard deviation of triplicate measurements from at least two independent experiments conducted in the presence of either NAD($^+$/H) or NADP($^+$/H). All the enzymatic activities were assayed in cell-free extracts obtained from exponentially-growing cells cultured on M9 minimal medium containing 20 mM glucose. In the case of activities represented by more than one enzyme, the cofactor specificity of the total activity is given.

### Description of other supplementary datasets

**Supplementary Data 1.** Raw $^{13}C$-labelling data. Raw intensities in counts for metabolites measured (0 = $M_0$ isotope, 1 = M + 1, etc.) in two biological replicates. Excel file.

**Supplementary Data 2.** Raw GC-MS data. GC-MS mass distribution vectors (MDV) were corrected for natural abundance from 20% [U-$^{13}C_6$]-glucose experiments. In the data, –15, –57, –85 and f309 represent the different fragments of derivatized amino acids with the respective fractional abundance for the isotopes $M_0$, M + 1, …, $M_{max}$. Relative flux ratios are averages from four biological replicates. Excel file.

### References


1. Winsor, G.L. *et al.* *Pseudomonas* Genome Database: improved comparative analysis and population genomics capability for *Pseudomonas* genomes. *Nucleic Acids Res.* **39**, D596-D600 (2011).
2. Winsor, G.L. *et al.* Enhanced annotations and features for comparing thousands of *Pseudomonas* genomes in the *Pseudomonas* Genome Database. *Nucleic Acids Res.* **44**, D646-D653 (2016).
3. Caspi, R. *et al.* The MetaCyc database of metabolic pathways and enzymes. *Nucleic Acids Res.* **46**, D633-D639 (2018).
4. Nikel, P.I., Martínez-García, E. & de Lorenzo, V. Biotechnological domestication of pseudomonads using synthetic biology. *Nat. Rev. Microbiol.* **12**, 368-379 (2014).
5. Nikel, P.I., Chavarría, M., Fuhrer, T., Sauer, U. & de Lorenzo, V. *Pseudomonas putida* KT2440 strain metabolizes glucose through a cycle formed by enzymes of the Entner-Doudoroff, Embden-Meyerhof-Parnas, and pentose phosphate pathways. *J. Biol. Chem.* **290**, 25920-25932 (2015).
6. Nogales, J. *et al.* High-quality genome-scale metabolic modelling of *Pseudomonas putida* highlights its broad metabolic capabilities. *Environ. Microbiol.* **22**, 255-269 (2020).
7. Nelson, K.E. *et al.* Complete genome sequence and comparative analysis of the metabolically versatile *Pseudomonas putida* KT2440. *Environ. Microbiol.* **4**, 799-808 (2002).
